# Developmental excitation-inhibition imbalance permanently reprograms autism-relevant social brain circuits

**DOI:** 10.1101/2025.09.07.674483

**Authors:** A Stuefer, G Colombo, S Gini, D Sastre-Yagüe, L Coletta, F Rocchi, M Aldrighetti, B D’Epifanio, G Como, L Balasco, N Barsotti, C Montani, FG Alvino, A Galbusera, S Migliarini, SM Bertozzi, M Pasqualetti, Y Bozzi, V Tucci, L Cancedda, MV Lombardo, A Gozzi

## Abstract

An influential theory proposes that an imbalance between excitation and inhibition (E:I) plays a central role in the etiology of autism and related developmental disorders. However, controversy exists as to whether this imbalance is a direct causal mechanism for autism, or a compensatory response to other primary etiological factors. Using chemogenetic manipulations in neonatal mice, we show that a transient E:I imbalance during development is sufficient to permanently reprogram autism-relevant social brain circuits. Chemogenetically manipulated mice exhibit lifelong impairments in sociability, persistent dysregulation of multiple autism-risk synaptic genes, and sustained cortical hyperexcitability in adulthood. Importantly, these social impairments are robustly rescued by pharmacological inhibition of neuronal excitability. Developmental E:I imbalance also disrupts functional connectivity in social brain regions enriched for transcriptionally dysregulated genes, suggesting a convergence of transcriptional and circuit-level pathology. Finally, multivariate modelling shows that behavioral dysfunction in chemogenetically manipulated animals closely associates with disrupted connectivity between prefrontal and mesolimbic dopaminergic regions. Collectively, our findings reconcile conflicting theories in the field and point to activity-dependent transcriptional remodeling as a foundational mechanism by which transient E:I imbalance during development can cause lasting, autism-relevant circuit dysfunction.

## Introduction

Autism spectrum disorder (ASD) affects approximately 1-2% of the population, making it one of the most common neurodevelopmental conditions^1, 2^. Autism is commonly thought of within the context of a ‘spectrum’ - a word that helps convey well-known and considerable heterogeneity that is multi-scale in nature (e.g., genome, neurobiology, clinical presentation, severity, etc.)^3^. For example, while genetic contributions are widely recognized as central to autism etiology^4^, the genetic landscape of the spectrum is remarkably heterogeneous^5^. Specifically, large-scale genetic studies have revealed that autism is associated with both highly penetrant, rare mutations (with more than 100 high-confidence autism-risk genes^6^), as well as multiple common genetic variants^4, 7^. Environmental factors, such as prenatal inflammation and immune dysfunction, have also been implicated in the etiology of autism^8^. Human and animal studies have shown that these etiological contributors affect diverse biological and developmental processes, including transcriptional regulation, synaptic activity, neurogenesis and cell migration^9, 10^.

The multitude of neurobiological processes associated with autism has prompted the question of how these heterogeneous factors may converge to produce the relatively narrow set of behavioral symptoms that characterize autism. In 2003, Rubenstein and Merzenich ^11^ proposed that a neurophysiological imbalance between excitation and inhibition (hereafter termed ‘E:I imbalance’) may serve as a unifying cross-etiological mechanism behind autism. According to this theory, abnormally increased excitation, or insufficient inhibition would cause cortical hyperexcitability, excessive spiking, and lower signal-to-noise processing in key neural circuits, ultimately leading to the autism phenotype. Most empirical support for this theory comes from genetic investigations showing that multiple high-confidence genetic variants associated with autism (e.g. *Tsc2*, *Shank3*, *Fmr1*, *CDKL-5*, *NLgn*) are critically implicated in the regulation of E:I balance, often through synaptic-related mechanisms^12, 13, 14^. Several mouse models harboring mutations in these genes exhibit heightened cortical excitability, spontaneous seizures or epileptiform activity^15, 16^. In keeping with these findings, human investigations have revealed that approximately one third of autistic individuals develop clinically apparent seizures^17^, with almost 50% of autistic children showing epileptiform activity in electroencephalography (EEG) or magnetoencephalography (MEG) recordings^18, 19^.

Although widely invoked, the causal role of E:I imbalance in autism etiology has recently been questioned, with three main lines of criticism challenging its foundational assumptions^12, 20, 21^. First, much of the supporting evidence for this theory derives from rare genetic mechanisms, whose generalizability to idiopathic autism remains unclear. Second, the direction of E:I changes remains heterogeneous, with evidence of both increased and decreased excitability in autism^22^. Finally, recent animal investigations suggest that E:I imbalance may not represent a primary cause of autism, but rather a homeostatic, compensatory response to other underlying etiological factors^21^. As a result, whether E:I imbalance is causative or compensatory in autism remains unclear.

These opposing views can be reconciled within a developmental perspective: transient alterations in the E:I ratio during critical developmental windows may alter brain development through activity-dependent transcriptional mechanisms, leading to maladaptive circuit assembly^23, 24^. Within this framework, developmental E:I imbalance would thus contribute to autism etiopathology primarily by derailing early circuit development, independent of the homeostatic cascade it may later produce. Here, we test this hypothesis using chemogenetic manipulations in the developing mouse neocortex. We report that transient neocortical E:I imbalance during an early postnatal window is sufficient to produce enduring autism-relevant multiomic phenotypes (i.e., transcriptomic, connectomic and phenomic). Notably, these alterations converge to disrupt socially-relevant circuits whose activity predicts impaired sociability and behavior in manipulated animals. Our results suggest that developmental E:I imbalance is sufficient to initiate a core etiopathological cascade of mechanistic relevance for autism.

## Results

### Chemogenetic activation of Vglut1-positive pyramidal neurons enables transient elevation of neocortical excitability during early development

To test the hypothesis that increased E:I balance during development is sufficient to produce autism-relevant phenotypes, we used a Cre-dependent genetic strategy to express the Gq-coupled DREADD receptor hM3Dq in Vglut1 positive pyramidal neurons. Because activation of hM3Dq by the exogenous ligand clozapine N-oxide (CNO) increases neuronal excitability and facilitates firing^25^, this approach allowed us to transiently elevate neocortical excitability within predefined developmental windows. We chose to express hM3Dq in Vglut1-positive neocortical neurons to model the broad hyperexcitability that may arise from mutations in autism-risk genes expressed across widespread cortical and hippocampal regions^14^. Vglut1-specific DREADD expression was achieved by crossing Vglut1-Cre mice with ROSA26-floxed hM3Dq mice. The resulting double transgenic mouse line is here referred to as Vglut1-gq.

Given our focus on the developmental effect of E:I imbalance, we first verified DREADD receptor expression in Vglut1-gq mice during the early postnatal period. Previous work showed that Vglut1 is detectable as early as postnatal day 0 in the rodent brain^26, 27, 28, 29^. Accordingly, we observed robust perinatal expression of hM3Dq in neocortical areas of Vglut1-gq pups (but not control Vglut1-cre mice) by post-natal day 5 (PND5, Supplementary Figure 1a-b). hM3Dq receptor expression remained elevated into the second postnatal week (Figure 1a) and was largely prominent in cortical areas and in the hippocampal formation (Figure 1a, Supplementary Figure 1a-c).

**Figure 1.**
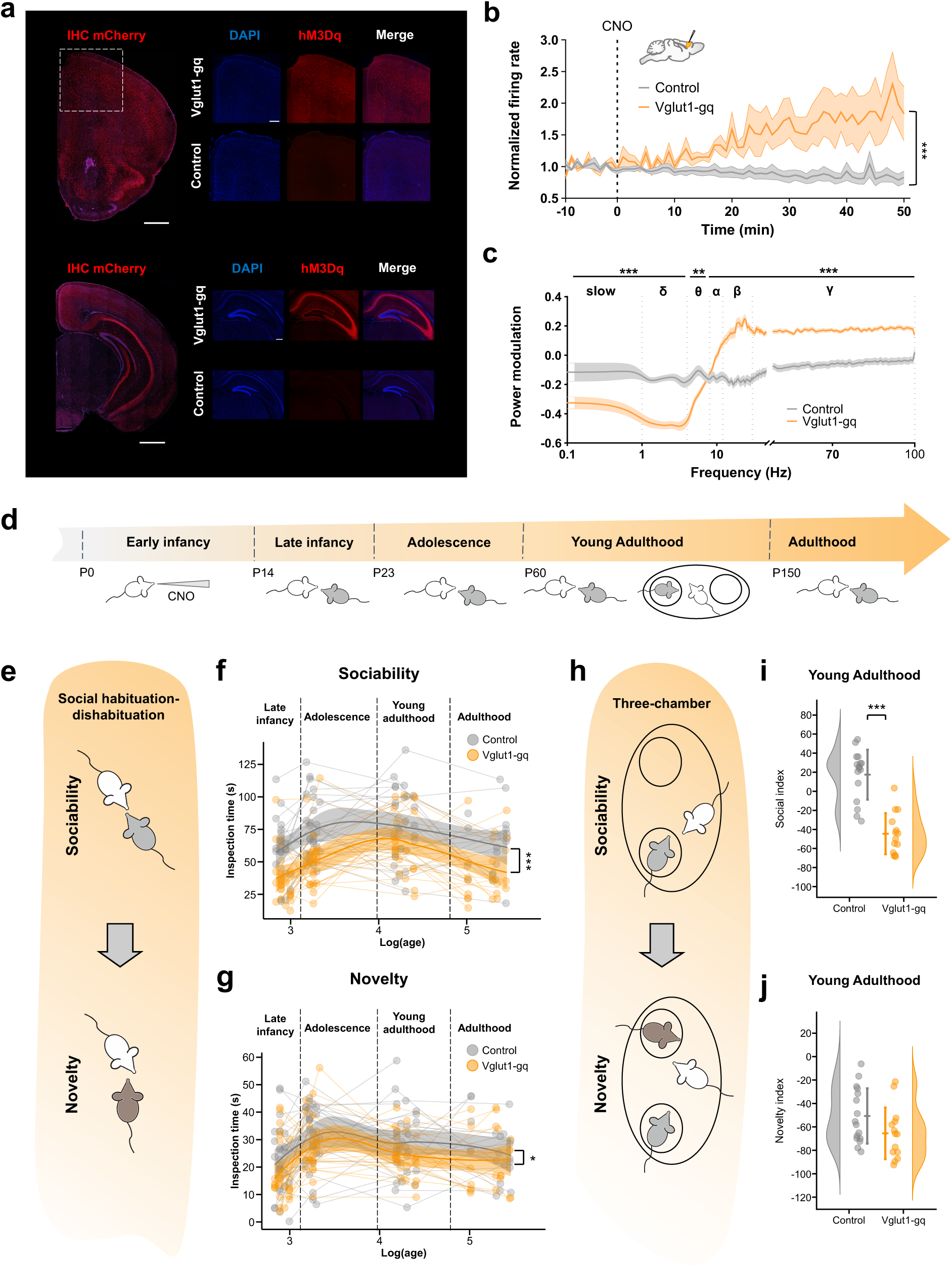
Developmental E:I imbalance leads to lasting impairments in sociability. **(a)** Immunohistochemistry showing hM3Dq (mCherry) expression and nuclear counterstaining (DAPI) in neocortex and hippocampus of control and Vglut1-gq mice at PND14. **(b)** Increased firing rate after CNO administration in Vglut1-gq mice (orange, n=4) compared to control mice (gray, n=5). Recordings were carried out on awake head-fixed mice (PND8-PND17). Two-way ANOVA, p<0.001. Data are presented as mean ± SEM. **(c)** Power spectral density of LFP in the PFC after CNO injection. Cluster-based permutation test, **p<0.01, ***p<0.001. **(d)** Experimental timeline. **(e)** Schematic of the social habituation–dishabituation task. **(f)** Vglut1-gq mice (n=26) exhibited a lasting reduction in sociability (inspection time) compared to control littermates (n=27; p<0.001). Shaded bars indicate mean ± 95% CI. **(g)** Vgluti1-gq mice showed reduced inspection time during novelty trial of the habituation dishabituation test (p=0.028). Shaded bars indicate mean ± 95% CI. **(h)** Schematic of the three-chamber test. **(i)** Reduced social index in Vglut1-gq mice (n=14) compared to control littermates (n=16; p<0.001.) **(j)** No difference in novelty index was observed between groups (p=0.120).

Having ascertained that DREADD receptors are perinatally expressed in Vglut1-gq mice, we next optimized a dosing regimen that enabled sustained CNO exposure in the neonatal brain. Oral administration of 5 mg/kg CNO resulted in elevated brain levels of CNO and its active metabolite clozapine for up to 6 hours post-administration (Supplementary Figure 1d-e). To confirm the efficacy of the employed dosing, we performed electrophysiological recordings in awake head-fixed pups (PND8-17) after administration of a pharmacokinetically equivalent subcutaneous CNO dose (1 mg/kg, Supplementary Figure 1d-e). CNO administration produced a robust increase in firing activity in the medial prefrontal cortex (PFC) of Vglut1-gq mice (Figure 1b, p<0.001) along with a marked reduction in slow band LFP power (slow, δ, θ, α band), and a concomitant broadband enhancement of high frequency (β and ƴ) LFP power (cluster-based permutation testing, q<0.01, all bands Figure 1c). These electrophysiological signatures have been consistently associated with increased cortical excitability^22^. Importantly, no overt epileptiform activity was observed upon CNO administration in any of our recordings. Taken together, these findings suggest that neonatal CNO administration in Vglut1-gq mice robustly increases cortical excitability, leading to an E:I imbalance.

### A transient E:I imbalance during early development permanently disrupts social behavior

To probe the behavioral impact of transient developmental E:I imbalance, we chronically administered CNO to Vglut1-gq and Vglut1-cre control littermates over the first two postnatal weeks. This postnatal period is developmentally comparable to late gestation and early infancy in humans, and encompasses multiple key developmental processes known to be altered in autism, such as neurogenesis, neural cell migration, synaptogenesis, and neural circuit formation^30^. The resulting cohorts (Vglut1-gq n=26 and control littermates n=27) underwent a comprehensive longitudinal behavioral assessment targeting multiple domains, including sociability, motor stereotypies, anxiety and working memory.

Notably, our behavioral investigations revealed that a transient developmental E:I imbalance leads to robust and persistent social deficits (Figure 1d-g, habituation phase, p<0.001, group effect, FDR corrected). Vglut1-gq mice exhibited marked social impairments (Figure 1f, Supplementary Figure 2), when tested during late infancy (p<0.001), adolescence (p<0.001), young adulthood (p=0.030) and adulthood (p=0.050). These deficits did not significantly differ across sexes (sex, p=0.092, group*sex, group*age, sex*age, group*sex*age, p>0.248, all interactions). Vglut1-gq mice also showed a mild reduction in time spent sniffing a novel conspecific during the novelty phase of the social habituation-dishabituation test (Figure 1g; p=0.028). However, this effect did not reach statistical significance at any individual age examined (Supplementary Figure 2c; all p>0.340).

We next conducted an additional set of control experiments aimed to probe the robustness and specificity of this social impairment. First, we asked whether this phenotype was replicable, and detectable using a different test of sociability. To this end, we conducted a three-chamber sociability test on an independent cohort of adult mice (Vglut1-gq n=14; control littermates n=16). In line with our initial findings, Vglut1-gq mice showed severely impaired sociability in this assay, exhibiting virtually no preference for the social stimulus over object exploration (Figure1h-i; p<0.001). No impairment in social novelty was observed in Vglut1-gq mice using this behavioral assay (Figure 1j, social novelty, p=0.121). Together, these results corroborate the notion that a transient increase in excitability during early development is sufficient to induce long-lasting social impairments.

To rule out the possibility that these alterations might be secondary to altered olfactory sensing or encoding, we conducted an olfactory habituation/dishabituation task on an independent cohort of Vglut1-gq and control mice, using both socially-neutral and socially relevant odors, at two different ages (late infancy, young adulthood). This test probes the ability of mice to detect and distinguish familiar and novel odors (water, almond, orange and 2 unfamiliar social odors^31^). Our measurements revealed preserved olfactory discrimination and memory in Vglut1-gq mice (Supplementary Figure 3). Specifically, the time spent sniffing both neutral and socially-relevant odors was comparable in Vglut1-gq and control littermates across trials (Supplementary Figure 3, water p=0.553, almond p=0.302, orange p=0.787, social 1 p=0.580, social 2 p=0.152) and ages (late infancy p>0.407, adulthood p>0.352, all stimuli). These results suggest that social impairment in Vglut1-gq mice is not epiphenomenal to the disruption of core olfactory functions.

### Impaired sociability in Vglut1-gq mice is not associated with overt sensory, motor or cognitive alterations

Our investigations show that developmental E:I imbalance produces lasting social impairments in Vglut1-gq mice. Are these deficits restricted to the social domain, or do they extend to other aspects of behavior? To address this question, we examined the impact of this manipulation on additional behavioral measures.

We first asked whether Vglut1-gq mice would exhibit other autism-relevant phenotypes by quantifying self-grooming, a proxy for repetitive and stereotyped behavior^32^. We observed a trend toward increased self-grooming that did not reach statistical significance (Supplementary Figure 4a-b p=0.091). In the same open field test, we observed no intergroup differences in total distance traveled (Supplementary Figure 4d-e, group, p=0.301), or in relative distance travelled in the center of the arena, an index of anxiety-like behavior^33^ (Supplementary Figure 4f, group p=0.362). Similarly, rotarod testing^34^ revealed no significant impairment in motor function and coordination in Vglut1-gq mice (Supplementary Figure 4c, group effect, time on the rotarod p=0.526, number of falls p=0.648). We next investigated the presence of tactile hypersensitivity using a whisker-nuisance test in adult mice. This assay probes the behavioral response to mechanical whisker stimulation, revealing heightened sensory responsivity in multiple genetic models of autism^35^. Responses to whisker stimulation were comparable between groups across all tested parameters (Supplementary Figure 5, p>0.492, all behaviors), suggesting unaltered tactile processing in the manipulated mice. Together, these findings indicate that the observed social impairment in Vglut1-gq mice is unlikely to reflect generalized deficits in motor, sensory, olfactory, or affective domains.

Because Vglut1-cre also drives hM3Dq expression in hippocampal regions, we next examined whether developmental E:I imbalance in Vglut1-gq mice would lead to hippocampal-dependent cognitive dysfunction. To this end, we used the Y-maze to assess working memory during exploratory behavior, and the novel object recognition task to assess long-term memory. In the Y-maze test we observed no intergroup differences in the number of spontaneous alternations of arm entry, number of entrances per arm or total arm entrances (Supplementary Figure 6a-d, group effect, p>0.261, all measures). Likewise, discrimination index scores, exploration time during habituation and test phase in the novel object recognition test were broadly comparable in Vglut1-gq and control mice (Supplementary Figure 6e-h; p>0.147). These results argue against the presence of major alterations in hippocampus-dependent cognitive function in Vglut1-gq mice and corroborate the domain specificity of the social phenotype induced by early E:I imbalance.

### A transient E:I imbalance during adolescence does not alter social behavior

To probe the developmental specificity of the observed social impairment, we chemogenetically increased excitatory activity in a separate cohort of Vglut1-gq mice during adolescence and subsequently performed behavioral testing in adulthood. The developmental window we probed (PND30–PND43) corresponds to puberty and adolescence in mice^36^. Interestingly, we observed preserved sociability in Vglut1-gq mice, with no changes in time spent sniffing during either habituation or novelty trials (Supplementary Figure 7a-d; sociability p=0.625; novelty p=0.911). We also observed no difference in anxiety-related behavior, locomotor activity (Supplementary Figure 7e; open field test, % distance traveled in the center, total distance traveled, p>0.769) or working memory in Vglut1-gq mice during adolescence (Supplementary Figure 7f; Y-maze, spontaneous alternations, number of entrances per arm or total arm entrances, p>0.614). These results corroborate the developmental specificity of our findings, suggesting that the social impairment observed in Vglut1-gq mice is specifically attributable to an E:I imbalance occurring in early development.

### Developmental E:I imbalance alters expression of autism-relevant synaptic genes

What developmental mechanisms could underlie the behavioral impairments observed in Vglut1-gq mice? Previous studies showed that increased neuronal activity can persistently alter gene expression through activity-dependent mechanisms^37^. We thus reasoned that transient hyperexcitability induced by CNO administration in Vglut1-gq mice could trigger similar adaptive responses, resulting in enduring transcriptional remodeling.

To test this hypothesis, we conducted RNA-seq on PFC samples collected in late infancy and adulthood from an independent cohort of Vglut1-gq and control mice (Figure 2a). This investigation revealed robust alterations in gene expression in Vglut1-gq mice, identifying 530 differentially expressed (DE) genes, of which 265 were upregulated (DE+), and 265 were downregulated (DE-; Figure 2a, group effect, FDR q<0.05 Supplementary Table 1). No significant group-by-age interaction was observed (n=0 DE genes, q>0.05), suggesting that developmental E:I imbalance can persistently remodel the transcriptional profile of affected brain regions.

**Figure 2.**
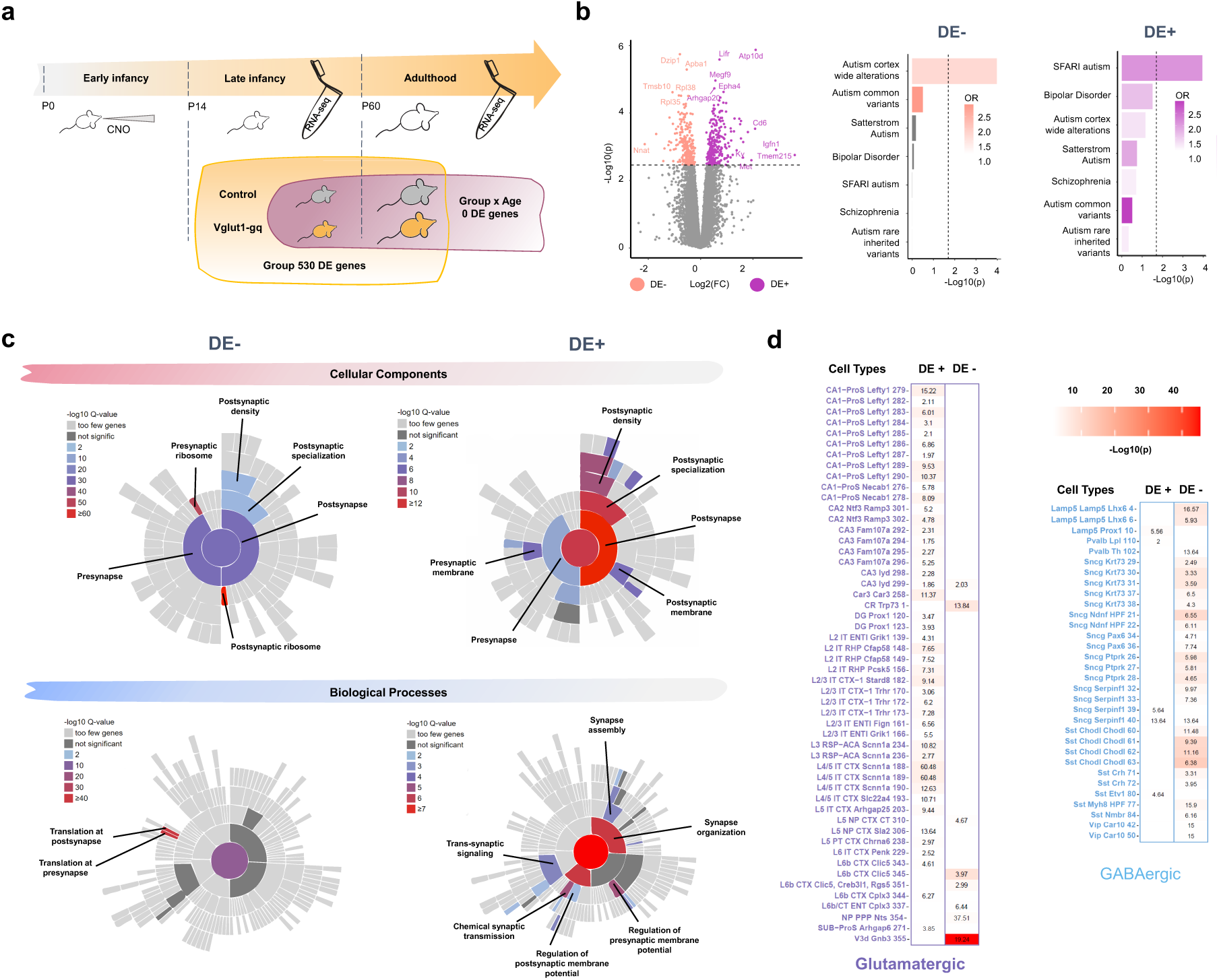
Developmental E:I imbalance alters expression of genes associated with autism and synaptic functioning. **(a)** Schematic of experimental timeline and summary of main results. **(b)** Volcano plot showing differentially expressed genes in Vglut1-gq mice (downregulated, DE–, salmon; upregulated, DE+, purple). DE– and DE+ sets show distinct enrichment patterns for autism-associated gene sets, but not for bipolar disorder or schizophrenia. Dotted line indicates FDR q<0.05. **(c)** Circular plot showing enrichment of DE– and DE+ genes for synaptic cellular components (top) and biological processes (bottom) as per the Syngo database. Color indicates enrichment significance as –log10 Q-value. **(d)** Enrichment analysis of DE+ and DE-gene sets with neural cell type markers from mouse single cell transcriptomic atlas of the Allen Institute. Numbers denote odds ratio (OR), color scale indicates -log10 p-value, as illustrated in the legend. Only significant enrichments at FDR q<0.05 are shown.

Given the postulated role of activity-dependent transcriptional mechanisms in the etiology of autism^38^, we asked whether the identified DE genes were significantly enriched for genes relevant to autism and neuropsychiatric disorders. This analysis revealed that both DE- and DE+ lists are robustly and specifically enriched for autism-relevant genes. In particular, the DE-list exhibited prominent enrichment for genes that are differentially expressed in the *post mortem* cortex of autistic individuals^39^ (Figure 2b, OR=1.706 q<0.001 Supplementary Table 2). The DE+ list was instead significantly enriched for known autism-risk genes belonging to the curated SFARI Gene database (https://gene.sfari.org; Figure 2b, OR=2.107 q<0.001 Supplementary Table 2), including syndromic autism genes like *GRIN2A*, *CDKL5*, *TCF4*, *KCNB1*, and *PHIP*. Notably, neither gene list revealed any significant association with genes associated with bipolar disorder or schizophrenia, thus suggesting that developmental E:I imbalance may primarily affect the expression of autism-related genes.

Given the critical role of activity-dependent transcriptional mechanisms in regulating synaptic function and their putative implication in the etiology of autism^38^, we examined whether the identified gene sets were enriched for synaptic-related genes. We found significant enrichment for both presynaptic and postsynaptic genes within the DE+ and DE− gene sets, with a prevalence of postsynaptic localization and a more robust synaptic involvement for the DE+ gene set (Figure 2c; FDR q<0.05). Further analyses revealed that the two gene sets were involved in partly dissociable biological processes: DE− genes exhibited robust enrichment for synaptic genes involved in protein translation, whereas DE+ genes showed enrichment for a broader set of synaptic processes, including synapse assembly and organization, regulation of synaptic membrane potential, neurotransmission and signaling (Figure 2c, FDR q<0.05 Supplementary Table 3).

We next performed an unbiased pathway enrichment analysis using the Reactome database to explore additional pathways of mechanistic interest within our DE gene sets (Supplementary Figure 8). In keeping with a possible activity-dependent origin for these transcriptional changes, we found that the DE-gene list was robustly enriched for genes involved in RNA metabolism (OR=29.371), autophagy (OR=7.847), DNA replication (OR=5.192), and chromatin organization (OR=7.338, FDR q<0.0497, all lists Supplementary Table 4). In contrast, no significant enrichment in additional non-neuronal ontologies was observed within the DE+ gene set.

To further interpret our DE gene sets within the context of relevant neuronal populations, we performed enrichment analyses against cell type-specific gene expression signatures. This investigation revealed a functional compartmentalization of transcriptional changes, with the DE+ set being predominantly associated with glutamatergic neuronal populations, and the DE-gene set being broadly enriched in markers of GABAergic neuronal populations (Figure 2d, Supplementary Table 5). These findings suggest that transcriptional remodeling produced by hyperexcitable pyramidal neurons may adaptively extend to inhibitory neuronal circuits. More broadly, our transcriptomic analyses indicate that early-life E:I imbalance can trigger reprogramming of synaptic gene expression, predominantly impacting autism-relevant pathways.

### Transient developmental E:I imbalance causes lasting cortical hyperexcitability

Given the altered expression of synapse-related genes and ion channels observed in Vglut1-gq mice, we asked whether transient developmental E:I imbalance would produce lasting changes in pyramidal neuron excitability. Whole-cell patch clamp electrophysiological recordings from L2/3 pyramidal neurons in the PFC of adult Vglut1-gq mice revealed broadly altered intrinsic electrophysiological properties in manipulated cells (Figure 3). Specifically, Vglut1-gq neurons displayed significantly lower input resistance, higher membrane capacitance, shorter membrane time constants, and increased rheobase compared with controls (Figure 3b, p<0.010 for all properties). Analysis of active firing properties revealed decreased firing rates at low current injections but increased firing at higher current injections (>300 pA, group X pA interaction p=0.001), as well as decreased first-to-last interspike interval ratio, representing lower firing adaptation (Figure 3c p=0.031). At the synaptic level, Vglut1-gq neurons exhibited significantly reduced sEPSC median amplitude (Figure 3d, p=0.025), but no significant changes in event frequency, decay kinetics, or charge transfer (p>0.418, all). These findings suggest that developmental hyperexcitability alters the intrinsic electrophysiological profile of the affected neuronal cells, resulting in nuanced input-dependent changes in neuronal responsiveness.

**Figure 3.**
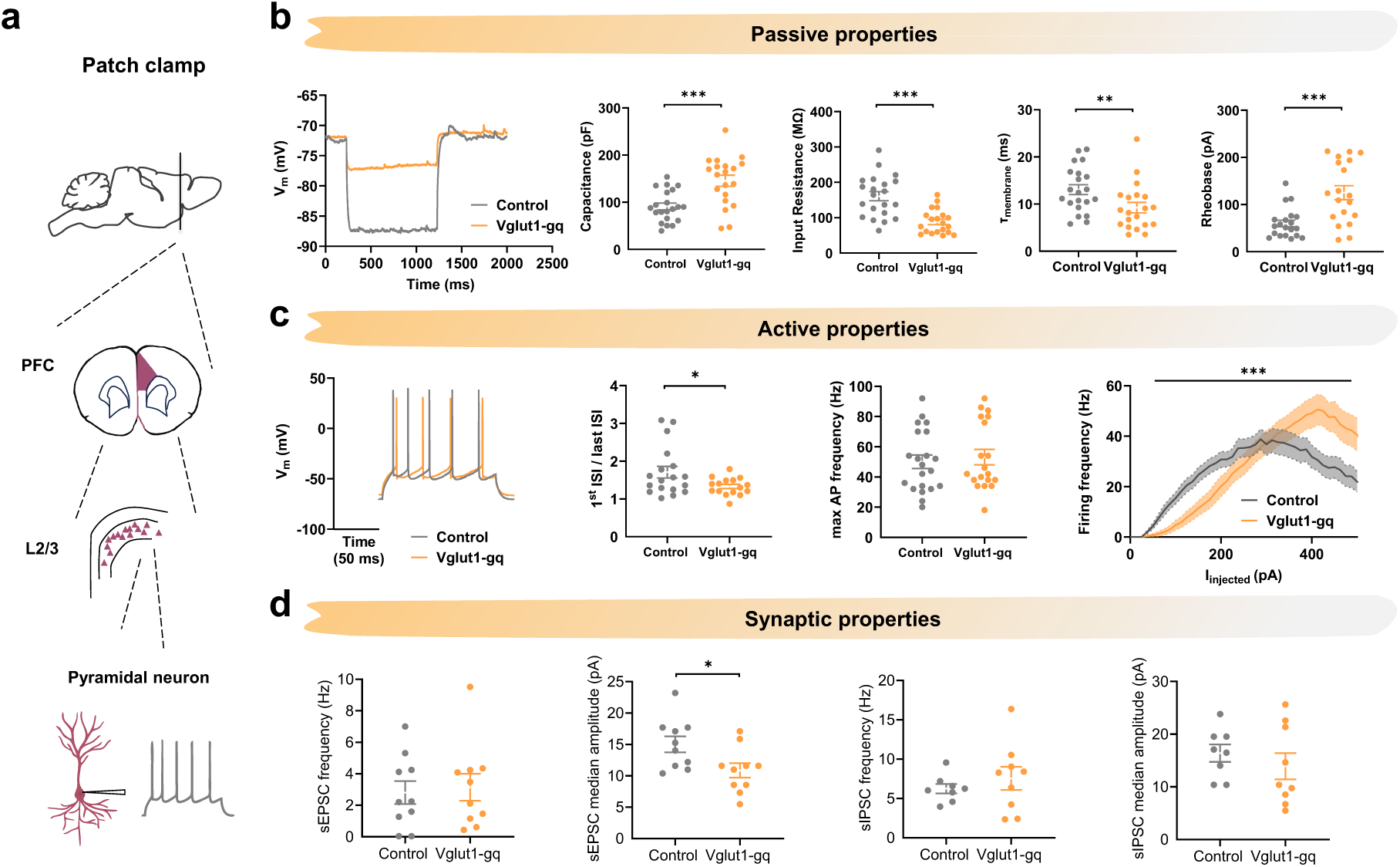
Developmental E:I imbalance remodels intrinsic properties of pyramidal neurons. **(a)** Schematic of whole-cell patch-clamp recordings from layer 2/3 pyramidal neurons in adult PFC (Vglut1-gq, orange, n=19 cells from 5 mice; control, grey, n=21 cells from 5 mice). **(b)** Passive properties. Representative V_m traces (left). **(c)** Active properties. Representative spike trains (left). Frequency–current (F – I) profiles show a significant group × current interaction (two-way RM ANOVA, p<0.001). **(d)** Synaptic properties (voltage clamp). Data are individual cells (dots), mean ± SEM. *p<0.05, **p<0.01, ***p<0.001, unpaired Student t test (unless otherwise stated).

To test whether these alterations persist at the network level, we performed EEG recordings in the PFC of adult Vglut1-gq and control mice (Figure 4a). Spectral analysis revealed a robust reduction in normalized EEG power across multiple frequency bands (δ, θ and α) in Vglut1-gq mice (Figure 4b, p<0.044). This EEG signature is consistent with increased cortical excitability in manipulated mice. Accordingly, the Hurst exponent (H) of EEG time series, an established marker of cortical excitability^22^, was significantly reduced in Vglut1-gq mice (Figure 4c, p=0.001). Together, these findings show that transient developmental E:I imbalance induces lasting alterations in population dynamics and cortical hyperexcitability.

**Figure 4.**
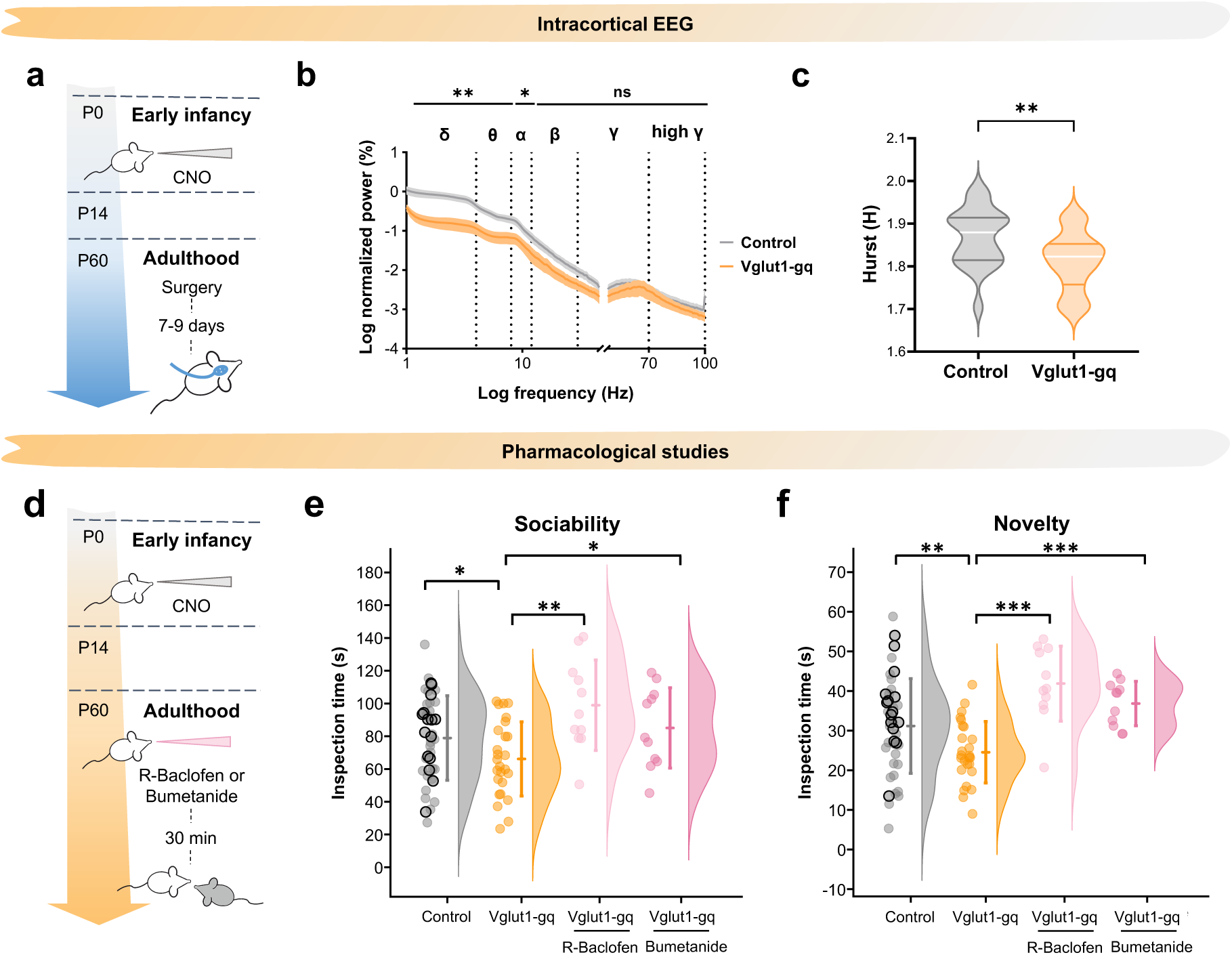
Developmental E:I imbalance leads to persistent cortical hyperexcitability in adulthood. **(a)** Schematic of the experiment. **(b)** Log-normalized power across frequency bands (δ 1–4 Hz, θ 4–8 Hz, α 8–12 Hz, β 12–30 Hz, γ 30–70 Hz, high-γ 70–100 Hz) in Vglut1-gq (orange; 21 recordings from 7 mice) and controls (grey; 21 recordings from 7 mice) *p<0.05, **p<0.01, ***p<0.001 Cluster-based permutation test. **(c)** Reduced Hurst exponent (H), in Vglut1-gq mice (**p<0.001). **(d)** Schematic of the pharmacological experiment. **(e)** Sociability (habituation trials) and **(f)** novelty (dishabituation trials) in control mice and Vglut1-gq mice treated with R-baclofen or bumetanide. Black-outlined grey circles denote control animals used in the new drug testing. Non-outlined grey circles represented control mice pooled from previous investigations. *p<0.05, **p<0.01, ***p<0.001, Student t test, FDR corrected.

### Pharmacological inhibition of cortical hyperexcitability rescues sociability in Vglut1-gq mice

Previous research has shown that altered cortical E:I balance results in impaired sociability in mice^40^. Given that Vglut1-gq mice exhibit persistent cortical hyperexcitability alongside impaired sociability, we asked whether acute reduction of excitability could restore social behavior. To address this question, we administered the GABA-B receptor agonist R-baclofen or the NKCC1 inhibitor bumetanide to adult Vglut1-gq and control mice prior to a social habituation/dishabituation test. Both compounds enhance GABAergic inhibition and have previously been shown to normalize neuronal hyperexcitability in mouse models of autism^41, 42^. Notably, both drugs effectively rescued social deficits in Vglut1-gq mice, restoring sociability to levels comparable to controls during both habituation and novelty trials (Figure 4d-f; p<0.041 and p<0.007, respectively). These findings show that pharmacological enhancement of GABAergic inhibition can acutely reverse social deficits induced by developmental E:I imbalance, implicating persistent cortical hyperexcitability as a mechanistic determinant of impaired sociability in Vglut1-gq mice.

### Developmental E:I imbalance selectively disrupts connectivity within the mouse social brain

What are the circuit-level bases of these behavioral alterations? Previous research has shown that synaptic-related alterations can profoundly affect large-scale interareal communication and functional connectivity in both autism mouse models and autistic individuals^43, 44^. We thus investigated the functional organization of large-scale brain networks in adult Vglut1-gq mice using resting-state fMRI.

We first asked whether cortico-cortical fMRI connectivity was broadly disrupted in Vglut1-gq mice. We thus assessed the functional integrity of the somatosensory network, one of the most resilient cortical systems in the mouse brain^45^. Consistent with unaltered sensory processing and motor responses in adult Vglut1-gq mice, fMRI connectivity within this network was largely preserved (Figure 5a-c, T>2). By contrast, seed-based analysis of the PFC, a key hub of the mouse default mode network (DMN)^46^, revealed robustly reduced fMRI connectivity in Vglut1-gq mice (Figure 5d-f). In line with the social impairments observed in these animals, connectivity disruption was particularly prominent between the PFC and hub components of the mammalian social brain, including orbitofrontal cortex, posterior cingulate cortex, and both dorsal and ventral striatum^47, 48^ (Figure 5e). Importantly, these connectivity deficits were not attributable to loss of excitatory or inhibitory neurons in the PFC, as postmortem quantification revealed comparable Vglut1- and Gad1-expressing cell numbers in Vglut1-gq and control mice (Supplementary Figure 9, p=0.651 and p=0.924, respectively). Collectively, these findings show that neocortical E:I imbalance during early development persistently and selectively disrupts the functional connectivity of socially-salient circuits.

**Figure 5.**
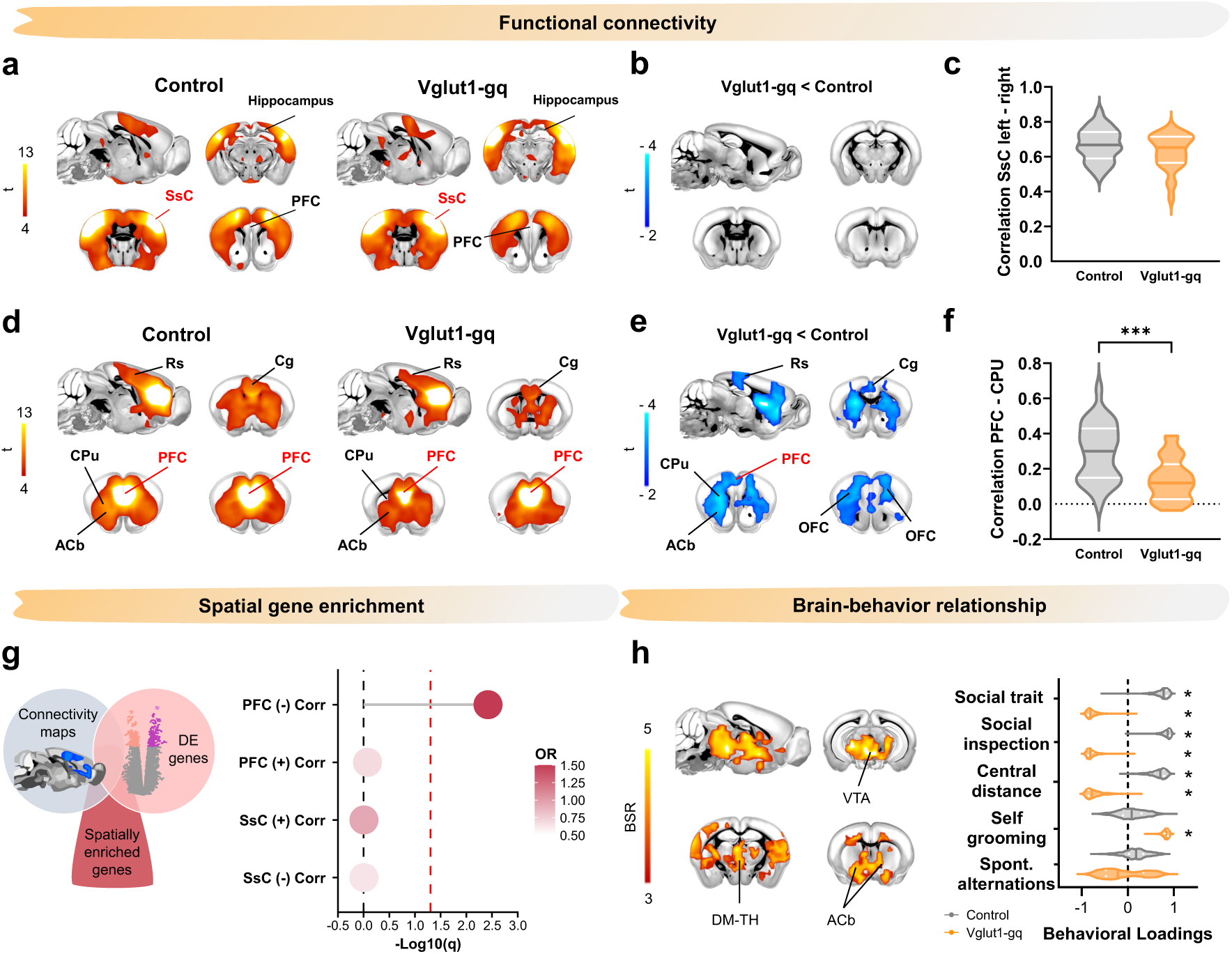
Developmental E:I imbalance disrupts connectivity and function of the mouse social brain. **(a)** fMRI connectivity map upon seed-based probing of the SsC in control (left) and Vglut1-gq mice (right) and **(b)** corresponding connectivity difference map. **(c)** Quantification of fMRI connectivity between left and right SsC. **(d)** fMRI connectivity map upon seed-based probing of the PFC in control (left) and Vglut1-gq mice (right) and corresponding **(e)** connectivity difference map. Blue indicates areas exhibiting significantly decreased fMRI connectivity in Vglut1-gq mice. **(f)** Quantification of fMRI connectivity between the PFC and Caudate Putamen. **(g)** Spatial gene–connectivity analysis. Left, schematic of the approach employed. Right; gene decoding and enrichment analyses illustrating significant overlap between DE genes and genes that are highly expressed in the mouse social brain (negative correlation OR=1.699, p=0.005). No such enrichment was observed for genes associated with somatosensory areas, in which fMRI connectivity is preserved (positive and negative correlations p=0.986, FDR corrected). **(h)** Behavioral PLS analysis. Bootstrap ratio (BSR) of brain loadings highlighting a mesolimbic dopamine network whose connectivity with PFC is predictive of behavioral performance in control animals. Red/yellow coloring represents bootstrap-ratio (BSR) z-scores obtained from 1000 bootstrapping iterations. This relationship is disrupted in Vglut1-gq mice (right). Columns indicate the contribution of each observed behavioral variable to the identified latent component. Asterisk indicates a significant contribution to the overall PLS relationship (95% CIs, not encompassing zero).

### Differentially expressed genes are spatially enriched within the mouse social brain

How could our broad neocortical manipulation preferentially disrupt connectivity within specific cortico-limbic regions while sparing others? We hypothesized that this effect may be explained by the spatial distribution of the DE genes identified in our study. Since these DE genes are vulnerable to transcriptional dysregulation induced by developmental E:I imbalance, we reasoned that they might be disproportionately expressed in brain regions critical for social behavior, and relatively less expressed elsewhere. Under this scenario, developmental E:I imbalance would primarily disrupt functional connectivity of regions and circuits anatomically enriched for these vulnerability transcripts.

To test this hypothesis, we asked whether DE genes were spatially enriched in regions exhibiting disrupted functional connectivity (Figure 5g). Using spatial gene decoding analysis from the Allen Mouse Brain Atlas, we identified N=757 genes whose pattern of expression spatially overlapped with the fMRI hypoconnectivity maps (Figure 5d-f). Among these, 669 genes showed higher expression in hypoconnected regions (“negatively correlated genes”), while the remaining 88 exhibited lower expression in these same regions (“positively correlated genes”). We reasoned that if developmental E:I imbalance preferentially dysregulates genes expressed within socially relevant regions, genes showing higher expression in hypoconnected areas (i.e., those negatively correlated) should be statistically enriched for DE genes. Consistent with this, enrichment analysis revealed a significant overlap between DE genes and negatively correlated decoded genes (OR=1.699, p=0.005), but not positively correlated decoded genes (OR=0.681, p=0.840; Figure 5g).

To probe the specificity of this result, we repeated the analysis using connectivity maps derived from primary somatosensory cortex, a region showing no connectivity alterations (Figure 5a-c). Consistent with our hypothesis, this control analysis revealed no significant gene enrichment (positively correlated n=43, OR=0.943; negatively correlated n=469 OR=0.628, p=0.986, FDR corrected, both measures; Figure 5g). Together, these results indicate that DE genes are preferentially expressed in brain regions relevant to social functioning, thus revealing a possible mechanism whereby developmental E:I imbalance can produce network-specific alterations in the mammalian brain.

### Prefrontal–dopaminergic circuit dysfunction is linked to social deficits in Vglut1-gq mice

Having established that developmental E:I imbalance disrupts prefrontal connectivity with cortico-limbic circuits, we next asked whether these alterations are behaviorally relevant. We reasoned that if functional dysconnectivity of these regions underlies behavioral alterations in Vglut1-gq mice, functional connectivity within these networks should be (a) closely associated with behavioral performance in control animals, and (b) this relationship should be disrupted in Vglut1-gq mice. We tested this hypothesis using a multivariate model (Partial least squares, PLS^49^) that identifies circuit components whose connectivity covaries with behavioral performance. Given the prominent role of prefrontal circuits in the behavioral alterations observed in Vglut1-gq mice, we modeled PFC connectivity against individual behavioral scores obtained from individual tests previously linked to the activity of this region^50, 51^. The use of aggregative behavioral measures in PLS has the advantage of reducing statistical bias thus minimizing risk of type-I error associated with repeated univariate testing^52^.

Corroborating the behavioral relevance of our connectivity findings, PLS analysis revealed one significant latent component (p=0.012) accounting for 44% of the covariance between PFC patterns of connectivity and behavioral performance. Notably, anatomical mapping of this component revealed a subcortical set of regions including the dorsomedial thalamus and key nodes of the mesolimbic dopamine system (i.e., ventral tegmental area, nucleus accumbens), whose connectivity robustly correlated with sociability and other behavioral traits in control mice (Figure 5h). This result supports the notion that the functional connectivity alterations we report here are behaviorally relevant. In keeping with this notion, we also found that this brain/behavior relationship was largely disrupted in Vglut1-gq mice (Figure 5h). Together, these results reveal a behaviorally relevant prefrontal–dopaminergic network whose connectivity predicts sociability and behavioral performance in control mice. Developmental E:I imbalance disrupts functional coupling within this network, thus implicating aberrant prefrontal–mesolimbic connectivity in the emergence of autism-relevant social dysfunction in mice.

## Discussion

Here we show that transient developmental E:I imbalance induced via chemogenetic activation of Vglut1-positive neocortical neurons is sufficient to produce multiomic, autism-relevant phenotypes in mice. These alterations encompass long-lasting transcriptional dysregulation of autism-related synaptic genes, cortical hyperexcitability, and robust deficits in social behavior. Additionally, we report selective disruption of functional connectivity within social brain circuits that are enriched for E:I vulnerability genes. Collectively, our results highlight a critical developmental window during which heightened cortical excitability can functionally reprogram the social brain, providing empirical mechanistic evidence that developmental E:I alterations might causally contribute to autism etiopathology.

The direct etiological relevance of E:I imbalance in autism has long been debated, with recent studies arguing that electrophysiological signatures of E:I disruption in autism mouse models may reflect compensatory adaptations to upstream etiological factors^21, 53^. Our work helps reconcile this controversy within a developmental framework. Specifically, our findings suggest that E:I imbalance during early critical windows may be only partially compensated, leading to maladaptive reprogramming of social brain circuits, plausibly through activity-dependent transcriptional mechanisms. The developmental specificity of our results, marked by the absence of social alterations in mice manipulated during adolescence, supports this framework and underscores its relevance for autism and early-onset developmental disorders.

Our findings corroborate and expand previous research on the long-term behavioral outcomes of developmental chemo/optogenetic manipulations in three important directions^54, 55^. First, we identify key transcriptional correlates of early-developmental perturbations, providing a mechanistic link between transient early-life hyperexcitability and lasting circuit and behavioral dysfunctions potentially relevant for autism. Second, we link developmental E:I imbalance to multiomic phenotypes with direct biological significance for autism, beyond the debated face-validity of autism-like social alterations in rodents. For example, the Hurst index reduction we found in EEG signals recorded in Vglut1-gq mice closely recapitulates analogous EEG signatures observed in autistic populations^22^. Similarly, our transcriptomic analyses revealed changes consistent with gene-expression alterations reported in postmortem cortical tissue from autistic individuals^39^. Likewise, fMRI-based functional hypoconnectivity within cortico-limbic circuits aligns closely with established neuroimaging findings in both genetic models of autism and human cohorts^44, 56^. These correspondences support the notion that the mechanisms we describe here are relevant for autism and related developmental disorders. Finally, our work reveals a plausible mechanism through which widespread developmental perturbations (in our case, a neocortical increase in E:I balance) can preferentially impact specific circuit substrates and behavioral domains relevant to autism. Our spatial gene decoding analyses suggest that these dysfunctions arise primarily in anatomical regions characterized by higher expression of genes vulnerable to (hyper)activity-dependent dysregulation.

While the observed cross-species correspondences support the translational validity of our findings for autism, these results should not, however, be interpreted as representative of the spectrum in its entirety. Rather, these phenotypes may help pinpoint the biological origins of specific subtypes within the broader autism spectrum, in line with the emerging view of autism as a biologically heterogeneous condition^3, 4, 57^. Within this framework, our experimental approach (and its extension to other experimental manipulations) may offer mechanistically relevant hypotheses regarding the etiological bases of subtype-specific neurobiological alterations identified with neuroimaging methods (e.g., fMRI, EEG) or via transcriptional analyses^22, 44, 58, 59^.

Our RNA-sequencing analysis revealed robust and lasting transcriptional alterations in Vglut1-gq mice involving numerous synaptic genes and key regulators of protein synthesis. The identity of these persistently altered transcripts and the nature of our manipulation support the notion that the observed transcriptional reprogramming is driven by activity-dependent mechanisms. This result aligns with accumulating literature showing that transient increases in neuronal activity can durably reshape gene expression via activity-dependent signaling pathways^23, 38^. Our results expand this framework by showing that transient neuronal hyperactivity during early development can lead to maladaptive reprogramming of this regulatory cascade, ultimately resulting in autism-relevant behavioral and network-level alterations.

Corroborating this link, the identified transcriptional signatures show strong convergence with molecular pathways implicated in autism, including synapse assembly, signaling, and protein translation, which we found to be both up- and downregulated in Vglut1-gq mice. Importantly, while loss-of-function mutations are frequently implicated in autism, dysregulation or duplication of autism-risk genes can equally produce developmental derailment and autism-related developmental syndromes^4, 60^. This notion is supported by multiple animal and human studies reporting autism-relevant phenotypes resulting from deletion or duplication of multiple genetic loci containing well-known autism-risk genes (e.g., *SHANK3*, *UBE3A*, *NRXN1*, *MECP2*, *SCN2A*, *TSC2*, *FMR1*^61, 62, 63, 64, 65, 66^). Our findings thus emphasize that balanced synaptic-related gene expression is essential for neurotypical development, and that deviations in this nuanced regulatory equilibrium may represent a key molecular mechanism underlying autism-related disorders.

Previous work has shown that E:I imbalance and altered synaptic signaling in the PFC can disrupt social behavior in mice^40, 67, 68^. Consistent with this framework, our EEG and pharmacological investigations converge to suggest that PFC hyperexcitability may underlie the social deficits produced by developmental E:I imbalance. Building on this finding, our neuroimaging studies also reveal some plausible network level correlates of impaired sociability in mice. Specifically, we found robustly disrupted connectivity between the PFC and several of its targets, including the posterior cingulate, dorsal striatum and regions of the mesolimbic systems. This finding is consistent with the result of acute chemogenetic studies showing that acute elevation of PFC excitability robustly disrupts fMRI connectivity and transiently impairs social behaviors in mice^69^. Our results also add to emerging evidence of an inverse relationship between fMRI connectivity and cortical excitability^44, 70^. Importantly, our PLS analysis highlighted a crucial role of PFC-mesolimbic and thalamo-prefrontal coupling in mouse sociability. This result suggests that social behavior critically relies on an effective top-down modulation of inputs converging onto reward-related circuits. More broadly, our findings provide a systems-level framework that reconciles the results of previous optogenetic studies showing a key contribution of multiple PFC-targeted regions (including the mediodorsal thalamus, retrosplenial cortex, and mesolimbic dopamine areas) in gating social behavior in rodents^71, 72, 73^.

In conclusion, our data show that neural hyperexcitability during a critical developmental window induces lasting alterations in autism-like social behavior, gene expression, cortical excitability, and functional connectivity, closely recapitulating hallmark phenotypes observed in autism. These findings provide direct empirical support for a causal role of developmental E:I imbalance in autism etiology, and offer key mechanistic insights into both the origin and significance of prevalent translationally-relevant autism phenotypes.

## Methods

### Ethical Statement

All in vivo experiments were conducted in accordance with Italian law (DL 26/214, EU 63/2010, Ministero della Sanità, Roma). Animal research protocols were reviewed and approved by the animal care committee of the University of Trento, Italian institute of technology and the Italian Ministry of Health.

### Animals

Experiments were conducted on double-transgenic animals of both sexes obtained by crossing R26-DIO-hM3Dq-mCherry^74^ with Vglut1-Cre (JAX stock #023527; Harris et al, 2014, DOI: <u>10.3389/fncir.2014.00076</u>) mice to generate R26-DIO-hM3Dq-mCherry::Vglut1-Cre referred to as Vglut1-gq. Vglut1-Cre and WT littermates were used as controls. Stimulus mice for social behavioral measures were sex and age matched C57BL/6J (C57) (Jackson Laboratories, strain No: 000664). Mice were group-housed under controlled temperature (21 ± 1 °C) and humidity (60 ± 10%) and maintained on a standard 12-hour light/dark cycle. Food and water were provided ad libitum.

### Immunohistochemical Analysis

On postnatal days 5 (PND5) and 14 (PND14), mice were deeply anesthetized, and perfused transcardially with 4% paraformaldehyde (PFA). Brains were post-fixed overnight at 4°C, and coronal sections (50 μm) were obtained using a vibratome (Leica). Brain sections were incubated for three consecutive nights at 4 °C with a primary rabbit anti-RFP antibody (ab62341, Abcam, 1:500) diluted in PBTriton (0.1 M PB, 0,3% Triton X-100, 5% horse serum). The following day, brain sections were washed three times with PBTriton, and further incubated overnight at 4 °C with VectaFluor Excel Amplified DyLight 594 Anti-Rabbit IgG secondary antibody (DK1594 Vector Laboratories). Samples were then rinsed in PBTriton and the amplification step was performed according to manufacturer’s instructions. The resulting sections were counterstained with DAPI and mounted onto slides. Images were acquired with Nikon AX confocal microscope using 10x or 20x Plan Apo objectives and the arge Image function of NIS Elements software.**Error! Reference source not found.**

### In Situ Hybridization Analysis

For *in situ* hybridization analysis fresh brain tissue was embedded in Tissue Tek (Sakura) and frozen at −80°C. Brains were then cryo-sectioned in the coronal plane (14 μM thick). VGlut1 and Gad1 riboprobe were retrotranscribed from cDNA amplified with the following primers: Vglut1 Forward primer CAGAGCCGGAGGAGATGA; VGlut1 Reverse primer TTCCCTCAGAAACGCTGG (NM_182993.1); GAD1 Forward primer TGTGCCCAAACTGGTCCT; GAD1 Reverse primer TGGCCGATGATTCTGGTT (NM_008077.6). ISH analysis was performed as previously described^75^ using digoxigenin-labeled VGlut1 and Gad1 antisense riboprobes (0.7 Kb and 1 Kb, respectively). A chromogenic reaction using NBT/BCIP (11681451001, Roche) was used as substrate for alkaline phosphatase. Images were acquired with a MacroFluo microscope (Leica) equipped with Nikon NIS-Elements software. For quantification, three coronal sections spanning the region of interest were obtained from each mouse (n=3 Vglut1-gq mice and n=3 control mice). Cell numbers were quantified and normalized to the section area to calculate cell density, resulting in a total of 9 datapoints per group.

### Pharmacokinetic Measurements

N=24 male C57BL/6J mice (age PND2) were used to probe CNS levels of CNO (CNO dihydrochloride, Hellobio) and its active metabolites upon oral or subcutaneous administration. Oral administration of CNO (dissolved in 5% sucrose solution) was performed using a 10 μl pipette tip, by gently placing the tip at the pup’s mouth. In s.c. administration studies, CNO was diluted in saline. Brains were collected one or six hours after CNO administration and homogenized in four volumes (v/w) of phosphate-buffered saline containing protease inhibitor (100:1). An aliquot of each brain homogenate was extracted (1:3) with cold CH₃CN containing 200 nM clozapine-d₄ as internal standard. Calibration curves were prepared in naïve mouse brain homogenates to cover the range from 1 nM to 10 μM. Three quality control samples were prepared by spiking the parent compound into naïve mouse brain homogenate to yield final concentrations of 20, 200, and 2000 nM. Calibrators and quality control samples were extracted (1:3) with the same extraction solution used for brain homogenates. All samples were centrifuged at 3,270 × g for 20 min at 4 °C, and the resulting supernatants were diluted (1:1) with water prior to LC-MS/MS analysis. Measurements were performed on a Waters ACQUITY UPLC-MS/MS system equipped with a triple quadrupole detector (TQD) mass spectrometer, an electrospray ionization (ESI) interface, and a photodiode array (PDA) detector (Waters Inc., Milford, MA, USA). Electrospray ionization was applied in positive mode. Compound-dependent parameters, including MRM transitions and collision energies, were optimized for each analyte. Chromatographic separation was achieved on an ACQUITY UPLC BEH C18 column (100 × 2.1 mm ID, 1.7 µm particle size) with a VanGuard BEH C18 pre-column (5 × 2.1 mm ID, 1.7 µm particle size) maintained at 40 °C, using water + 0.1% formic acid (A) and acetonitrile + 0.1% formic acid (B) as mobile phases. Levels of the parent compound in brain tissue were determined from MRM peak area response factors and expressed as amount per gram of brain tissue.

### Chemogenetic Manipulations

Our pharmacokinetic measurements showed that oral administration of CNO at 5mg/kg (Supplementary Figure 1d-e) resulted in brain concentration levels sufficient to pharmacologically engage hM3Dq receptors^76, 77^ and comparable to those obtained with s.c. administration of CNO at 1 mg/kg, a treatment regimen widely used in previous studies^78, 79, 80^. On this basis, chemogenetically-induced developmental E:I imbalance was induced via daily oral administration of CNO (5 mg/kg) for 13 consecutive days, starting at PND1. The drug was dissolved in a 5% sucrose solution and administered at a volume of 1 μl/g to both Vglut1-gq and Vglut1-cre littermates. Drug treatments were consistently performed at the same time of day to minimize potential circadian influences. During the dosing period, animals were monitored daily for body weight gain and developmental milestones, such as the day of eye opening and the day of fur appearance.

CNO administration during adolescence (PND30-43) was carried out subcutaneously. For these studies, CNO (0.5 mg/kg) was dissolved in saline, and administered to both Vglut1-gq and control littermates. This regimen was selected based on pilot studies where activation of hM3Dq receptors in PFC CamkIIα neurons produced robust reductions of sociability and increased neuronal firing, without signs of epileptiform activity^69^. Comparable CNO doses have been widely used to elicit DREADD-induced neural activation^74, 81^.

### In Vivo Electrophysiology

#### Headpost Surgery

To perform head-fixed electrophysiological recordings in awake mouse pups (PND14), head-post implantation surgeries were conducted on n=4 Vglut1-gq and n=5 Vglut1-cre control littermates. Pups were anesthetized with isoflurane (5% for induction, 2% for maintenance), placed on a heating pad to maintain a body temperature of 37 °C, and secured in a stereotaxic frame (Stoelting Co.). The skull was exposed, lightly scored, and a small hole was drilled above the prefrontal cortex (PFC) for electrode insertion. A head-post (Neurotar) was affixed to the skull in two steps: first, dental cement (Superbond C&B kit, Dental Leader) was applied and the head-post gently pressed into place until set; then, a second cement (Paladur, Kulzer S.r.l.) was applied to further stabilize the implant. After surgery, pups were placed in a resting cage for 30 min to recover from anesthesia.

#### Electrophysiological Recordings

For acute electrophysiological recordings, CNO was administered via a s.c. cannula at 1 mg/kg. S.c. administration was employed to minimize motion and avoid potential interference with the recording apparatus that could occur with oral delivery. This dose yields brain concentrations of CNO and clozapine comparable to those produced by oral administration of 5 mg/kg, i.e. the regimen used in our developmental manipulations (p>0.81, all comparisons; Supplementary Figure 1d-e).

After surgery mice were allowed to recover from anesthesia for one hour. Following recovery, pups were head-fixed in the electrophysiology setup, and a s.c. cannula was inserted in the intrascapular region for CNO administration. A single-shank electrode (Cambridge Neurotech) was then advanced through the dura mater at a rate of 1 μm/min using a microdrive system (Kopf Instruments, Germany). Once the electrode reached the PFC, the recordings started. Neural activity was recorded for 20 min (baseline) in consecutive 5-min bins. CNO was then administered via the s.c. cannula, and recording continued for a further 50 min. Signals were amplified with an RHD-2000 system (Intan Technologies) and digitized at 20 kHz using the RHD Recording Controller Software (v2.09).

#### Local Field Potential and Multi-Unit Activity

MUA was calculated as described in Rocchi et al. (2022)^70^. Briefly, a 4th order Butterworth filter (<100 Hz cutoff) was applied to obtain high-frequency component of the signal. Data were then band-passed filtered between 400 and 3000 Hz using a Kaiser window filter (transition band: 50 Hz; stopband attenuation: 60 dB; passband ripple: 0.01 dB). Spike times were detected from this high-frequency signal using a threshold corresponding to 4 times the median of the high-frequency signal, divided by 0.6745 as described in Quiroga et al. (2004)^82^. Firing rate was computed for each electrode as the number of spikes divided by the recording duration in seconds (spikes/s). This rate was segmented into 1-min bins and averaged across channels. For normalization, firing rate in each channel was divided by the mean firing rate during the 10-min baseline window immediately preceding CNO injection (−10 to 0 min). Normalized traces were then averaged across channels for each animal, and group-level time courses were obtained by averaging across animals.

LFP signals were computed as described in Rocchi et al. (2022)^70^. Briefly, raw extracellular recordings were first down-sampled to 4 kHz and then band-pass-filtered to 1-250 Hz by applying a two-step procedure^83, 84^. First, a 4th order Butterworth filter with a cutoff frequency of 1 kHz was used to low-pass filter the time series. Data were next downsampled to 2 kHz, and further filtered with a Kaiser window filter (0.1-250 Hz, sharp transition bandwidth 0.1 Hz, passband ripple: 0.01 dB, stop band attenuation: 60 dB). Finally, the signals was resampled at 1 kHz. All filtering was performed in both the forward and backward temporal directions.

Power spectral density of LFP signal was computed in Matlab using the pspectrum function with a frequency resolution of 1 Hz. Power spectra were pooled over 1-min bins and then averaged across channels. Frequency-resolved spectral profiles were obtained by averaging the power spectra over time. A modulation index was then computed for each frequency as: (post-CNO channel-averaged power – baseline channel-averaged power)/(post-CNO channel-averaged power + baseline channel-averaged power). Group-averaged modulation index traces (± SEM across animals) were plotted for each experimental group. Frequency bands (slow (0.1-1 Hz), delta δ (1-4 Hz), theta θ (4-8 Hz), alpha α (8-12 Hz), beta β (12-30 Hz), and gamma ƴ (30-100 Hz)) were indicated with vertical dotted lines and used for statistical comparisons

Frequency resolved statistical quantifications were performed using a cluster-based permutation as described in Maris and Oostenveld ^85^. Briefly, a two-sided t-test was computed at each frequency, and adjacent significant frequencies (p<0.05) were grouped into clusters (C_i_). Cluster-level statistics were obtained by summing t-values within each cluster (T*i0* = ∑*f∈Ci*t*0*(f)). A permutation distribution was generated by shuffling group levels 104 times and recomputing the maximum cluster statistic for each permutation (T^p^). Corrected p-values were derived by comparing observed cluster statistics (T_i_^0^) to this permutation distribution (T^p^), as follows:

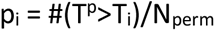

Significant clusters (p<0.05) were then assigned to their corresponding canonical frequency bands, and significance was indicated in the figure with asterisks above the relevant band labels.

### Behavioral Testing

#### Experimental Cohorts

Behavioral tests were carried out on different cohorts of animals. The primary dataset consisted of a large cohort of mixed-sex Vglut1-gq mice (n=26; 13 females, 13 males) and control littermates (n=27; 12 females, 15 males) that underwent the following behavioral assays: open field, social habituation/dishabituation, Y-maze, whisker nuisance, rotarod, and novel object recognition. Open field, social habituation/dishabituation and Y-Maze task were carried out longitudinally at three timepoints, late infancy (PND16-21), adolescence (PND23-29) and young adulthood (PND64-83). fMRI connectivity mapping in adulthood (PND62–80) was also performed in this original cohort of animals, to which we added three additional subjects (Vglut1-gq n=1, male; controls n=2, 1 female and 1 male) that did not take part in behavioral testing, but were amenable to fMRI imaging. Additional testing of social habituation/dishabituation was carried out in adulthood (PND100–259) on a smaller subset of this initial cohort of animals(consisting of Vglut1-gq, n=22; control, n=20). This choice reflected the exclusion of some animals due to attrition over the course of the study.

Grooming behavior was scored manually from video recordings of the open field test during minutes 10–20 of the session. Due to suboptimal video quality in several recordings, reliable scoring was possible only for 22 Vglut1-gq and 20 control littermates out of the main experimental cohort. In young adulthood (PND64–83), a subset of randomly selected animals (n=23 Vglut1-gq and n=20 control littermates) was further evaluated in the whisker nuisance, rotarod, and novel object recognition paradigms.

Two independent cohorts of animals were used for the olfactory habituation/dishabituation (PND15-18 and PND63-68, n=14; 9 female, 5 male Vglut1-gq mice and n=15; 7 female and 8 male control littermates) and 3-chamber sociability test, respectively (n=14; 10 female and 4 male Vglut1-gq mice and n=16; 6 female and 10 male control littermates).

Mice that received chronic CNO treatment during adolescence (PND30–43) were tested in adulthood (PND58–68) on the open field, social habituation/dishabituation, and Y-maze tasks (n=26; 11 female and 15 male Vglut1-gq mice; n=26; 13 female and 13 male control littermates).

All mice were acclimated to the test room for a minimum of 30 minutes before all behavioral tests. All testing sessions were filmed using a USB day and night camera (Ugo Basile), and animal position was continuously recorded by a video tracking system (ANY-Maze V6, Stoelting).

#### Social Habituation/Dishabituation Test

Mice underwent the social habituation/dishabituation test as described by Ferguson et al. (2000)^86^. Briefly, each mouse was individually placed in a testing cage (Tecniplast GR900, lightly illuminated at 6 ± 2 lux) one hour prior to the test. A stranger mouse of the same sex and age was introduced into the testing case for one minute allowing free interaction, and then removed for 3 min before being reintroduced. This sequence was repeated for four habituation trials with the same stimulus mouse. In the fifth trial (dishabituation or novelty trial), a new unfamiliar stranger mouse of the same sex and age was introduced for a one-minute interaction. Social interactions (termed “social inspection” here) was manually scored as time spent sniffing and/or following the stimulus mouse. Sociability was computed as the sum of social inspection during the first three habituation trials, whereas social novelty was quantified as social inspection during the sole dishabituation phase.

#### 3-Chamber Sociability Test

The 3-chamber sociability test was conducted in adult mice as described by Ferretti et al. (2019)^87^. Briefly, each mouse was placed in the center chamber of the apparatus with the doors to the side chambers closed for 10 min. The doors were then opened, allowing free exploration of all chambers for 10 min. In the subsequent sociability phase (10 min), wire cups were placed in the lateral chambers. One cup contained a stimulus mouse and the other remained empty. In the social novelty phase (10 min), a novel stimulus mouse was placed in the previously empty wire cage. Stimulus mice were age- and sex-matched C57 mice. Sniffing of the stimulus mouse was scored manually, and the social and novelty indices were calculated as described by Rein et al. (2020)^88^;

Social index = (time spent sniffing stimulus mouse-time spent sniffing empty wire cage) / total time spent sniffing × 100

Novelty index = (time spent sniffing familiar stimulus mouse-time spent sniffing novel stimulus mouse) / total time spent sniffing × 100

#### Olfactory Habituation/Dishabituation Test

The olfactory habituation/dishabituation test was performed in late infancy and adulthood as described by Yang and Crawley (2009)^31^. Five different odors were presented in three consecutive 2-min trials each, with a 1-min intertrial interval. The odors included water, almond (Paneangeli, 1:100 dilution), orange (Paneangeli, 1:100 dilution), and two unfamiliar social odors (cage swabs of sex-matched group-housed mice). Mice were acclimated to the testing cage (Tecniplast GM500, lightly illuminated at 6 ± 2 lux) for 30 minutes. After acclimation, cotton swabs with different odors were presented, and time spent sniffing was manually scored.

#### Rotarod Test

The rotarod test^34^ consisted of two habituation days followed by one test day in which the animal was placed on a rotating beam at varying speeds (rotations per minute, rpm). Adult mice were habituated with 5-min sessions at 4 rpm. On the test day, each mouse performed three 5-min trials, separated by 5-min intertrial intervals, with the rotation speed gradually increasing from 4 to 64 rpm. Trials ended upon the mouse’s fall or when the animal completed three full passive rotations on the beam. Time on the rotarod, rpm per minute, and number of falls were recorded as measures of motor performance for group comparisons.

#### Open Field Test

Mice were placed in the center of the open field arena (square apparatus 60 x 60 x 30 cm with black walls and a grey floor) and allowed to freely explore for the entire test duration (30 min)^33^. In the recording software (ANY-maze v6, Stoelting), a center (30 × 30 cm) and peripheral zone (remaining area) were defined, and ANY-maze was used to automatically compute the total distance traveled, as well as the distance traveled and the time spent in the center zone.

#### Grooming Behavior

Grooming behavior^89^ was assessed by manually scoring self-grooming from video recordings of the open field test. Scoring was performed over a 10-min window from minute 10 to 20 of the session, with the first 10 min of recording regarded as an acclimation period to the testing arena.

#### Whisker Nuisance Test

Adult mice underwent the whisker nuisance task as described by Balasco et al. (2019)^35^. Two days before the test, mice were allowed to acclimate to the testing cage (Tecniplast GM500) for 30 minutes. On testing day, animals underwent four sessions of five minutes each, including one sham session followed by three stimulation sessions. During the sham session, a wooden stick was introduced into the cage without contacting the mouse. During stimulation sessions, whiskers were stimulated by deflecting vibrissae with a wooden stick, and six behavioral responses (evading, guarding, freezing, attacking, climbing, and startling) were manually scored. The percentage of time spent evading, guarding, and freezing, as well as the number of attacking, climbing, and startling events, were computed.

#### Y-Maze Test

The Y-Maze test was performed as described by Faizi et al. (2011)^90^. Mice were placed in the center of a Y-shaped maze (three symmetrical gray arms at a 120-degree angle, 26 × 10 x 15 cm, illuminated at 20 ± 2 lux) and allowed to freely explore for 8 minutes. An arm entry was counted when all four limbs of the mouse were within an arm, and three consecutive entries into different arms were defined as a triad. Spontaneous alternations were computed as (number of triads/[total arm entries −2] × 100).

#### Novel Object Recognition Test

The novel object recognition test was carried out in adult mice and consisted of habituation and object acquisition on day 1, followed by the novel object recognition phase 24 h later^91^. During habituation, mice freely explored the testing arena (60 × 60 × 30 cm; black walls, grey floor) for 15 min. In the acquisition phase, two identical objects were placed in the arena for an additional 15 min of free exploration. After 24 h, each mouse was placed in the same arena with one familiar object from the acquisition phase and one novel object for 15 min. Object exploration was scored manually, and the time spent exploring each object during both the acquisition and the novel object recognition sessions was recorded. The discrimination index was calculated as (time spent investigating the novel object - time spent investigating the familiar object) / total time spent investigating objects.

#### Statistical Analysis

Behavioral tests were analyzed using a linear mixed-effects model in R-studio (v. 4.2.2). Fixed effects included group (Vglut1-gq vs. control), sex, age (for longitudinal tests), and their interactions. Random effects included random intercepts for dam and individual mouse identity (slope for age within each mouse for longitudinal measures). In tests where repeated trials within a session were the relevant source of variation (e.g., rotarod, whisker nuisance), trial was included as a fixed factor in place of age. *Post hoc* analyses at individual time points were corrected for multiple comparisons using the false discovery rate (FDR) procedure.

### RNA Sequencing

#### Tissue Collection and RNA Extraction

Tissue collection and RNA extraction were carried out as described by Montani et al. (2022)^92^. Briefly, Vglut1-gq and control mice were euthanized by cervical dislocation, their brains were extracted and placed in a mouse brain matrix (Agnthos, Sweden). Two 1-mm coronal sections at the level of the PFC were collected using scalpel blades. Sections were placed on semi-frozen glass slides, and 0.5-mm micropunches were used to isolate PFC tissue from each hemisphere and section. PFC samples were immediately frozen on dry ice and stored at −80 °C until RNA extraction.

All procedures were performed under RNase-free conditions. For RNA extraction, samples were disrupted and homogenized with motor-driven grinders for 3 min, and total RNA was isolated using the RNeasy Mini and Micro Kits (Qiagen). RNA concentration was measured with the Qubit RNA BR Assay Kit (Life Technologies). Purity was assessed by UV absorbance ratios (260/280 and 260/230) using a NanoDrop ND-1000 spectrophotometer (Thermo Fisher Scientific). RNA quality was determined by RNA integrity number (RIN) using the Agilent RNA 6000 Nano Kit on an Agilent 2100 Bioanalyzer (Agilent Technologies, Santa Clara, CA, USA), following the manufacturer’s instructions. All samples had RIN values > 8.

#### RNA Library Preparation and Sequencing

Library preparation was performed using the TruSeq® Stranded mRNA Sample Preparation Kit (Illumina, San Diego, CA, USA) with 500 ng input RNA per sample. All libraries were prepared in a single batch and sequenced with the NovaSeq 6000 S2 Reagent Kit (200 cycles) to an average depth of 100 million paired-end reads. Libraries (0.85 nM in 150 μl) were loaded onto an Illumina NovaSeq 6000 System (IIT–Center for Human Technologies, Genomic Unit, Genoa, Italy). Reads were aligned to the mm10 genome (GRCm38 primary assembly, Gencode) using STAR, and raw read counts were generated with the *featureCounts* function in R using Gencode v24 annotations. Sequencing-related variables were quantified using Picard tools functions (https://broadinstitute.github.io/picard/).

#### Analysis

RNA sequencing analysis was conducted using custom-written scripts in R (R-Studio version 4.2.2). Low-read genes, with less than 100 reads in two or more samples, as well as genes with low variance (variance filtering at 15%), were excluded from the analysis, resulting in a total of 11,819 retained genes for further analysis. Library size normalization was performed using the trimmed mean of M values (TMM) method^93^ with the calcNormFactors function of the edgeR R-library. Next, data were transformed to log-counts per million using the *voom* function of the limma package in R^94^ and precision weights were estimated for incorporation into linear modeling of differential expression. To account for sequencing-related artifacts, principal component analysis (PCA) was performed on the Picard variables (percent coding bases, percent utr bases, percent intronic bases, percent intergenic bases, median CV coverage, median 5’ to 3’ bias, aligned reads, and AT dropout) and further correlated with principal components (PCs) of the expression data. This analysis showed that sequencing-related artifacts were not correlated with the first PCs of the expression data and, therefore, did not need to be accounted for in further analyses.

DE genes were identified using functions for linear modeling in the limma library in R. The employed linear model included the main effects of group, age, and sex, as well as their interactions and RIN as a covariate (∼ group*age*sex + RIN). Multiple testing correction was performed using the false discovery rate (FDR q< 0.05).

Enrichment analysis was performed with the enrichR library in R (https://maayanlab.cloud/Enrichr/)^95, 96^. A series of curated gene lists of relevance for multiple neuropsychiatric disorders (Simons Foundation Autism Research Initiative (SFARI https://gene.sfari.org-downloaded April 2023), Gandal et al. (2018; 2022)^39, 97^,Satterstrom et al. (2020)^6^, Wilfert et al. (2021)^98^, Grove et al. (2019)^99^, the Allen cell type database (2021), the Reactome pathway database and the SynGO platform^100^ were employed. Custom code was used to compute enrichment odds ratios (OR) and p-values based on the hypergeometric distribution. A background total equivalent to the total number of genes analyzed after filtering low-read genes and low-variance genes (i.e., 11,819) was used, and gene identities (IDs) were converted from mouse gene IDs to human gene homologs. FDR correction (FDR<0.05) was applied.

### Patch Clamp Recordings

Patch-clamp experiments were performed in young adult mice (PND47-56). Animals were anesthetized with isoflurane and transcardially perfused with ice-cold artificial cerebrospinal fluid (ACSF) containing 117 mM NaCl, 3.5 mM KCl, 1.2 mM NaH₂PO₄, 2 mM MgCl₂, 1 mM CaCl₂, 25 mM NaHCO₃, and 25 mM glucose (∼310 mOsm, pH 7.4; oxygenated with 95% O₂ and 5% CO₂). For all experiments, brains were removed and immersed in cutting solution containing 220 mM sucrose, 5 mM MgCl₂, 2 mM MgSO₄, 2.5 mM KCl, 1.2 mM NaH₂PO₄, 0.5 mM CaCl₂, 25 mM NaHCO₃, 10 mM D-glucose, 10 mM Na-ascorbate, and 3 mM Na-pyruvate (∼300 mOsm, pH 7.4; oxygenated with 95% O₂ and 5% CO₂). 270 mm-thick coronal slices (cut with VT1000S Leica Microsystem vibratome) were allowed to recover for 30 minutes at 35°C in a solution containing: 117 mM NaCl, 3.5 mM KCl, 1.2 mM NaH₂PO₄, 3 mM MgSO₄, 0.5 mM CaCl₂, 25 mM NaHCO₃, 25 mM glucose, 5 mM Na-ascorbate, 3 mM Na-pyruvate, and 3 mM myo-inositol (∼310 mOsm, pH 7.4; oxygenated with 95% O₂ and 5% CO₂). All recordings were performed at room temperature in ACSF. Patch pipettes (3–5 MΩ) were pulled from thick-walled borosilicate glass capillaries (Sutter Instrument, catalog #B150-86-7.5) and filled with intracellular solution containing (in mM): 130 K-gluconate, 10 HEPES, 7 KCl, 0.6 EGTA, 4 MgATP, 0.3 NaGTP, and 10 phosphocreatine. The pH was adjusted to 7.3 with HCl.

Pyramidal neurons in layers II/III of the PFC were identified by their stereotypical orientation (perpendicular to the cortical surface) and characteristic pyramidal morphology. Firing properties and action potential (AP) shape were confirmed after break-in. Resting membrane potential (Vm) was measured in I=0 mode immediately after establishing whole-cell configuration. AP number was quantified by injecting current pulses from −50 to 520 pA (500 ms duration, 10 pA increments). The minimum current required to elicit the first AP was defined as the threshold current. To calculate rheobase, additional current pulses (500 ms duration, 1 pA increments) were applied starting from the threshold.

To calculate input resistance (Rin), 20 consecutive current steps (−50 pA start, +5 pA increments, 1 s duration) were injected, and the resulting steady-state membrane voltage deflection (ΔV) was recorded. Rin was then calculated as the ratio ΔV/I, where ΔV is the steady-state change in membrane voltage and I is the corresponding injected current. Membrane capacitance (Cm) was also calculated from the same 10 voltage responses to hyperpolarization current steps. All traces were averaged, and the decay phase of the voltage response was fit with a single exponential function. The mean time constant was then used to calculate Cm. Spontaneous and miniature excitatory postsynaptic currents (sEPSCs and mEPSCs) were recorded for 5 min in voltage-clamp mode at the resting membrane potential of the recorded cells. Recordings of sEPSCs and mEPSCs were obtained after 5 min of bath perfusion with 10 μM bicuculline or 10 μM bicuculline plus 1 μM tetrodotoxin (TTX), respectively (HelloBio, catalog #HB0893 and #HB1035). Data from neurons with access resistance >25 MΩ were discarded. Access resistance was continuously monitored by applying a 10-mV hyperpolarizing step (1000 ms) between protocols.

### EEG Recordings

#### Surgery

Mice were anesthetized with isoflurane inhalation (2–5%), secured in a bite bar, and placed on a stereotaxic apparatus (model 930; Kopf, CA). Artificial tear gel was applied to the eyes to prevent corneal drying. Once the mouse was deeply anesthetized, a midline sagittal incision was made along the scalp to expose the skull. l. A Foredom dental drill was used to create 1-mm holes over the right frontal lobe (+3.0 mm, −1.1 mm relative to Bregma, anterior– posterior axis). Two-channel electrode posts (Winslow, 267-7400) were anchored to 1-mm stainless steel screws (Bilaney Consultants GmbH, 00-96×3/32). Screws were advanced into drilled holes until secure, taking care not to penetrate the dura. An additional screw was placed on the left hemisphere to ensure proper stability of the implant. Dental cement was applied around the screws, at the base of the post, and on the exposed skull. At the end of surgery, mice received subcutaneous injections of ketorolac (5–7.5 mg/kg) and Baytril (5 mg/kg) for analgesic, anti-inflammatory, and antibiotic treatment. Body weight was monitored for three days post-surgery, and supplementary injections of ketorolac or Baytril were administered as appropriate. Mice were then individually housed, returned to the vivarium and monitored daily until EEG recordings (7-9 days after surgery day).

#### EEG Recordings

EEG signals were obtained using a Data Science International^TM^ (DSI) system from awake and freely moving mice (n=7 Vglut1-gq mice; 4 females, 3 males; n=7 control littermates; 3 females, 4 males). Mice were habituated to the recording environment for 15 min before connection to the DSI acquisition system. A two-channel tether was connected to the mouse’s two-channel electrode post (implanted during surgery) under brief isoflurane anesthesia. Mice were then placed back into recording apparatus after recovery. The tether was connected to a commutator positioned above the cage, and animals were further habituated for 15 min before recording. EEG signals were amplified with the 7700 Conditioner (DCOM) module and acquired with the ACQ-770 interface (PNM-P3P-7002XS, DSI). Signal quality was monitored online using Ponemah software (v5.50, DSI). Acquisition settings included a high-pass filter (>0.5 Hz), a low-pass filter (<100 Hz), and a sampling rate of 1 kHz. Each mouse underwent three 10-min recording sessions on the same day (total n=42 sessions).

#### EEG Data Analysis

Single EEG recordings were exported in European Data Format (EDF) and analyzed using custom MATLAB scripts (MathWorks, R2019b). LFP signals were preprocessed and analyzed following previously described methods^70^. Briefly, raw EEG signals sampled at 1 kHz were low-pass filtered with a finite impulse response (FIR) filter designed using a Kaiser window (passband: 249 Hz; stopband: 250 Hz; passband ripple: 0.01 dB; stopband attenuation: 60 dB) to remove high-frequency noise. Zero-phase filtering (filtfilt) was applied to avoid phase distortion. For each channel and recording, power spectral density (PSD) was estimated with the *pspectrum* function at a frequency resolution of 0.8 Hz. Spectra were normalized to the total power within 1–150 Hz and expressed as percentage power per frequency bin. Frequency bands were defined as: δ (1–4 Hz), θ (4–8 Hz), α (8–12 Hz), β (12–30 Hz), ϒ (30–70 Hz), and high-ϒ (70–150 Hz). For each band, mean relative power was extracted for group comparisons.

Differences between groups were assessed using a permutation-based test at the mouse level, preserving repeated recordings. Group labels were shuffled 10,000 times for each frequency band, and two-sided p-values were derived by comparing the observed difference to this distribution. This approach preserves repeated measurements within each mouse and controls for variability across recordings.

Single EEG recordings were further analyzed to estimate the Hurst exponent, a measure of long-range temporal correlations in the EEG signal^22^. Custom MATLAB scripts (MathWorks, R2019b) were used for preprocessing and Hurst exponent computation. Briefly, signals were band-pass filtered between 1–80 Hz using a third-order Butterworth filter (butter) and zero-phase filtered with filtfilt to avoid phase distortion. The filtered Local Field Potential (LFP) signals were then segmented into non-overlapping 30-second windows (corresponding to 7,500 samples at a sampling rate of 250 Hz). For each 30-second window and recording channel, the Hurst exponent was estimated using a multifractal detrended fluctuation analysis approach implemented in the bfn_mfin_ml function. Parameters for the multifractal analysis were set as follows: Haar wavelet filter (filter=’haar’), lower bounds (lb=[-0.5, 0]), upper bounds (ub=[1.5, 10]), and verbosity enabled for diagnostic output. Group-level comparisons of Hurst exponent values were performed using linear mixed-effects (LME) models implemented in MATLAB’s fitlme function. Fixed effects included sex, group (Vglut1-gq vs. control), and their interaction, while random intercepts were included for recordings nested within each mouse (H_mean ∼ sex * group + (1 | mouse_id) + (1 | mouse_id:recording_id)).

### Pharmacological Rescue of Social Impairment

A pharmacological rescue experiment was conducted in a separate cohort of Vglut1-gq mice (n=11; 8 females, 3 males) and control littermates (n=6; 5 females, 1 male). For analysis, we included Vglut1-gq mice from previous experiments. Additionally, control mice from this cohort were pooled with previously collected control groups to reduce animal use and increase statistical power as per the recommendation of the ethical review panel. All mice received CNO treatment during the first two postnatal weeks as described for other cohorts. Drugs (R-Baclofen, 1 mg/kg, Tocris Bioscience, or Bumetanide, 0.2 mg/kg, Merck Life Sciences B3023) were dissolved in saline and administered 30 min prior to social habituation/dishabituation testing in adulthood (> PND60).

The social habituation/dishabituation task was performed as described above. Briefly, animals were isolated for one hour before testing. During this isolation period mice were injected i.p. with either R-Baclofen or bumetanide. Treatment order was counterbalanced, with half of the mice receiving R-Baclofen in the first testing session and bumetanide in the second, and the remaining half receiving the reverse order. Testing sessions were conducted with an inter-session interval ranging from several days to several weeks. Following isolation, a same-sex, age-matched conspecific was introduced for a one-minute interaction, followed by a three-minute interval. This was repeated for four habituation trials with the same stimulus mouse, followed by a dishabituation trial introducing a novel unfamiliar conspecific. Social behaviors, including sniffing and following, were manually scored, and total inspection time was quantified during the first three habituation trials and the novelty trial. Statistical differences were assessed using t-tests with subsequent FDR correction for multiple testing.

### Functional Magnetic Resonance Imaging

#### Image Acquisition

Resting state fMRI (rsfMRI) was performed in adult mice. rsfMRI scanning was carried out with a light sedation protocol combining microdosing of chlorprothixene (Cpx 0.15 mg/Kg intramuscular injection) and isoflurane (0.5% administered via intubation). Cpx microdosing has been previously employed in functional studies to stabilize neural recordings without compromising cortical responsiveness or inducing burst suppression^101, 102, 103, 104^. Pilot studies in our lab showed that Cpx+IF produced functional networks closely comparable to those obtained with standard sedation mixtures (Pepe et al., in preparation). A key advantage of the employed protocol is that obviates the need for intravenous infusion, a factor that greatly simplifies operations and minimizes attrition in longitudinal studies.

Animal preparation was performed as previously described^70, 105, 106^. Briefly, mice were anesthetized with isoflurane (5% induction, 2% maintenance), followed by an intramuscular injection of Cpx. Animals were then intubated, artificially ventilated, and the isoflurane concentration was reduced to 0.5%. A 30-min stabilization period was allowed before the start of acquisitions.

rsfMRI image acquisition was performed on with a 7 Tesla MRI scanner (Bruker, Ettlingen) using a 72 mm birdcage transmit coil and a 3-channel solenoid coil for signal reception). The scanner was operated with Paravision 6.01 software (Bruker, Ettlingen). BOLD rsfMRI time series (1,920 volumes; 32 min) were acquired with an echo-planar imaging (EPI) sequence using the following parameters: repetition time (TR)/echo time (TE), 1,000/15 ms; flip angle, 60°; matrix, 98 × 98; field of view (FOV), 2.3 × 2.3 cm; 18 coronal slices; slice thickness, 550 µm; bandwidth, 250 kHz.

#### Image Preprocessing and Analysis

Image preprocessing and analysis were performed as previously described^70, 105, 106^. The first 50 volumes of each time series were discarded, and data were despiked, motion-corrected, skull-stripped, and spatially registered to a common reference template for adult mice. Motion traces (3 translations, 3 rotations) and average BOLD signal within a ventricular mask were used as nuisance covariates and regressed out from the time series. Data were then band-pass filtered (0.01–0.1 Hz) and spatially smoothed with a 0.6-mm full width at half maximum (FWHM) kernel. Volume censoring was applied using a framewise displacement (FD) threshold of 0.05 mm^107^. Seed-based functional connectivity was computed with custom seeds for the PFC, and the SsC for all subjects. The PFC seed was designed to span the entire antero-posterior extension of the prelimbic cortex, i.e. a hub component of the mouse PFC^108^. Group differences were assessed with two-tailed two-sample t-tests (t>2, p<0.05), corrected for multiple comparisons with family-wise error (FWE) cluster correction at p=0.05.

### Gene Decoding and Enrichment Analysis

To probe the potential mechanism underlying the predominantly social effects of the developmental E:I imbalance, we set out to test the hypothesis that the genes impacted by our manipulation (i.e., the differentially expressed (DE) genes), are preferentially expressed in areas of disrupted fMRI connectivity, which overlap with those involved in social behavior. To test this hypothesis, we implemented gene decoding^109^ and enrichment analyses of our fMRI connectivity difference maps. The goal of these analyses is to identify genes whose expression is spatially correlated with the topography of our “functional dysconnectivity” map (i.e., the unthresholded, between-groups connectivity difference seed-based maps obtained from prefrontal regions – social brain - or somatomotor cortex – control network). To this aim, we first obtained gene expression data for 75 brain regions defined in Rubinov et al. (2015)^110^ using *abagen*^111^. Among these, 68 regions had both gene expression and fMRI connectivity data. For each of these regions, we next assessed the correlation between gene expression levels and the magnitude of functional dysconnectivity using a linear regression. The resulting p-values were corrected for multiple comparisons, and genes with FDR-corrected p<0.05 (hereafter ‘decoded genes’) were retained for further analysis. This yielded 757 spatially enriched genes for the PFC and 512 for the somatosensory cortex map.

Gene enrichment analyses were then performed to assess whether there is a statistically significant overlap between the genes that are typically more expressed in the areas that display altered connectivity (i.e., the decoded genes) and those genes affected by the induced developmental E:I imbalance (i.e., the DE genes). Such an overlap would suggest that genes epigenetically impacted by our manipulation are also spatially enriched in regions exhibiting hypoconnectivity. The resulting p-values were further corrected for multiple comparisons within each map, using FDR correction.

### Partial Least Square Correlation Analysis

Behavioral Partial Least Square Correlation (PLS) was computed as previously described^49, 112, 113^. PLS is a data-driven multivariate approach that identifies weighted linear combinations (latent components) of two sets of variables that maximize shared covariance. Here, we performed a grouped behavioral PLS to identify patterns of covariance between functional connectivity and behavioral measures that were shared across Vglut1-gq and control animals, irrespective of absolute group differences. To this end, brain and behavioral data were normalized within each group, and a separate covariance matrix (R) was computed for each. We modeled PFC connectivity with behavioral measures from the same animals, all of which underwent the social habituation/dishabituation test (social trait, social inspection), open field test (central distance, self-grooming), and Y-Maze task (spontaneous alternations).

Behavioral measures were selected according to two criteria: (i) alignment with the test battery performed in the same cohort of mice that also underwent fMRI, and (ii) inclusion of the main readout of each task, with the addition of measures encompassing autism-related phenotypes. For the social habituation/dishabituation task, we focused on sociability, the primary outcome of this paradigm. We included both a single-age measure of social inspection in adulthood as well as a lifetime summary statistic of sociability across multiple ages (social trait), reflecting our interest in sociability as the core phenotype. For the open field test, we used central distance as the primary measure of anxiety-related exploration, complemented by self-grooming, which was included given its relevance to repetitive behaviors in autism models. For the Y-Maze task, the primary outcome of spontaneous alternations was used as a measure of working memory.

The R matrices from control and Vglut1-gq groups were then concatenated and subjected to singular value decomposition (SVD). Because sex was treated as a covariate of no interest, its effect was regressed out from connectivity maps prior to PLS^112^. The null hypothesis that the observed brain-behavior relationship could be due to chance was tested via permutation testing (1000 iterations) of the behavioral data. To prevent LCs from being driven by intergroup differences, permutation was performed within groups. The stability of brain and behavioral loadings was estimated with bootstrapping (1,000 iterations), with resampling performed within groups^112^.

## Acknowledgments

This work has been funded by the Simons Foundation (SFARI; grant number 982347 to A.G, M.V.L.), European Research Council (ERC) under the European Union’s Horizon 2020 research and innovation program ((#DISCONN; no. 802371 and no. 101125054 #BRAINAMICS to A.G.), and by the Brain and Machines Flagship Programme of the Italian Institute of Technology. A.G. is also supported by an endowment by Paolo and Sara Baracchino. M.V.L. acknowledges funding by the ERC under the European Union’s Horizon 2020 research and innovation program under grant agreement no. 755816. M.P. was supported by the EU H2020 MSCA ITN Project “Serotonin and Beyond” (no. 953327); the Next Generation EU National Recovery and Resilience Plan and Ministry of University and Research (no. ECS 00000017 “Tuscany Health Ecosystem THE,” Spoke 8); MIUR, Grant of the Department of Excellence 2023–2027; and MIUR PRIN 2022 PNRR (P2022ZEMZF). Y.B. was supported by TRAIN – Trentino Autism Initiative. University of Trento Strategic Project Research Grant 2018-2022. The authors are also grateful to Giuliano Iurilli and Francesco Papaleo for their generous feedback and inspiring discussions.

## Conflict of Interest

Cancedda, L. is funder and scientific consultant @IAMA therapeutics. Authors declare no further conflict of interest.

---

**Supplementary Figure 1.**
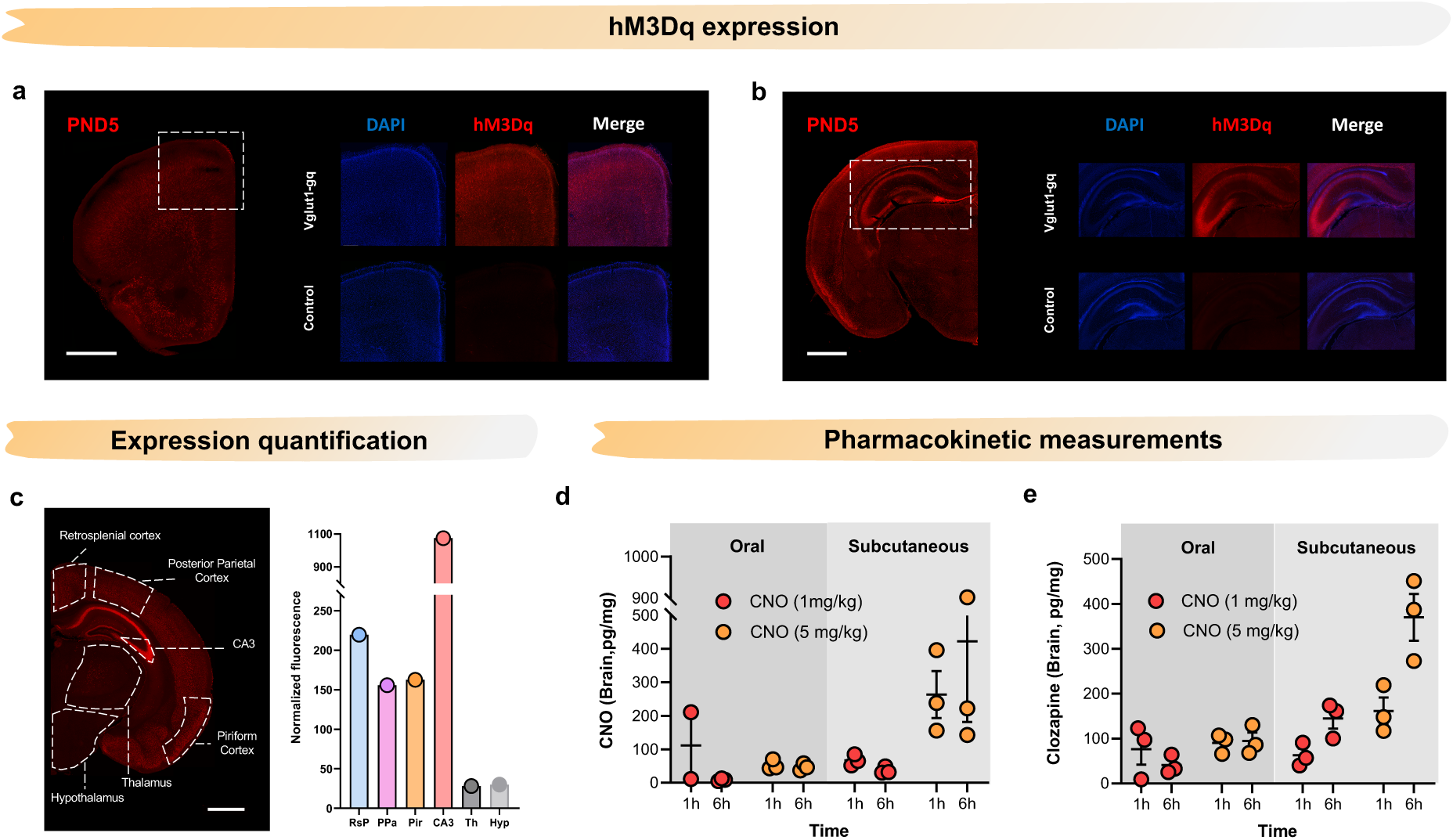
Expression of hM3Dq and pharmacokinetic measurements. **(a-b)** Immunohistochemical staining of hM3Dq at PND5 (DAPI, blue, hM3Dq-mCherry-red). **(c)** hM3Dq fluorescence quantification in representative regions of interest at PND14, (Retrosplenial cortex, RsP; Posterior parietal cortex, PPa; Piriform cortex, Pir; Thalamus, Th; Hypothalamus, Hyp). **(d)** Pharmacokinetic measurements of brain levels of CNO and **(e)** clozapine (pg/mg) in mouse pups 1 hour or 6 hours after oral (dark grey) or s.c. (light grey) CNO administration. Error bars indicate ± SEM.

**Supplementary Figure 2.**
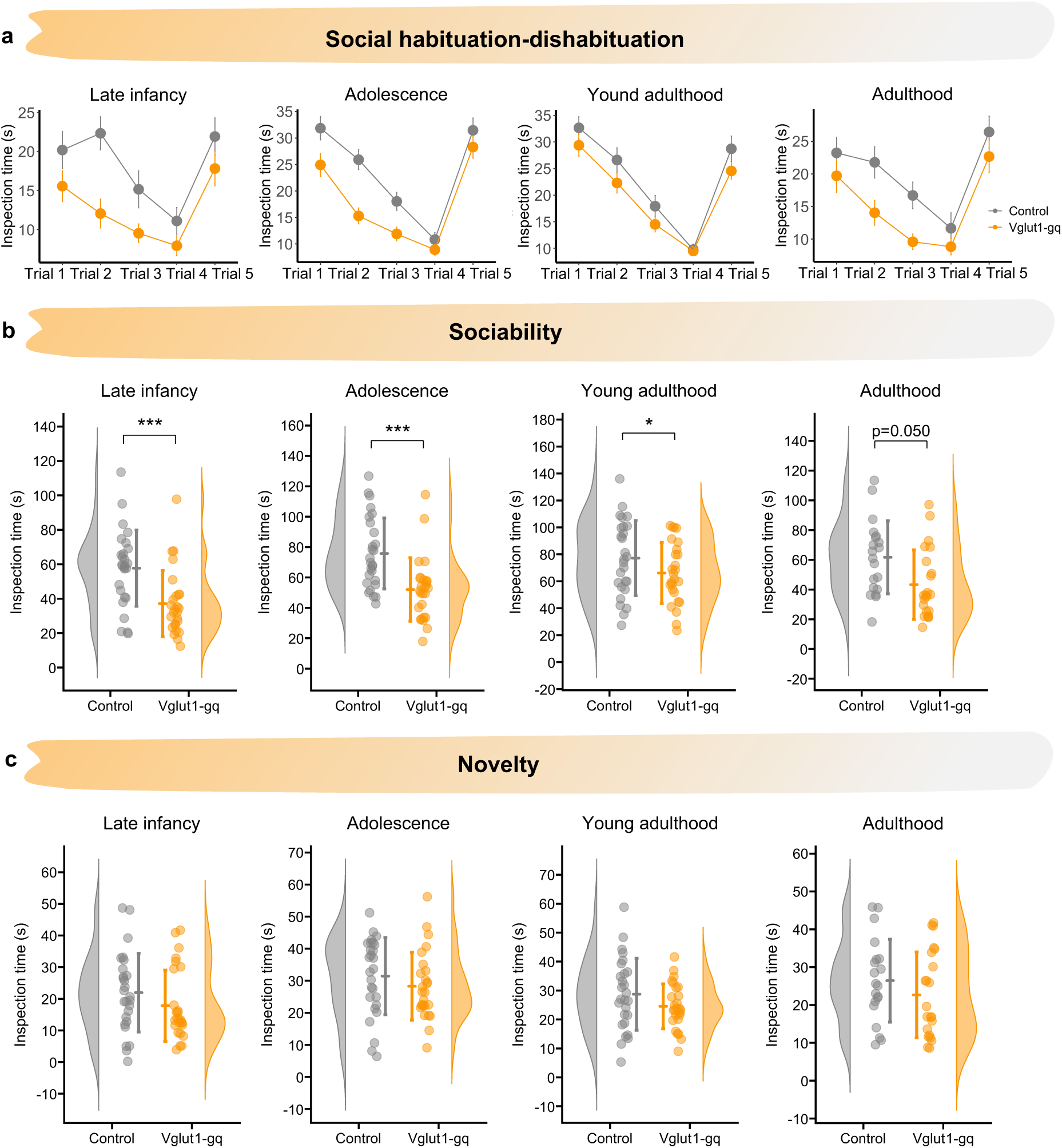
Social habituation-dishabituation test across development. **(a)** Inspection time across all individual trials at all the tested ages (late infancy, adolescence young adulthood Vglut1-gq, n=26, control, n=27; adulthood Vglut1-gq, n=22, control, n=20). **(b)** Cumulative inspection time (trials 1-3) during habituation (*p<0.05, p**< 0.01, ***p<0.001, FDR corrected). **(c)** Inspection time during the novelty trial across ages (p>0.340 all ages).

**Supplementary Figure 3.**
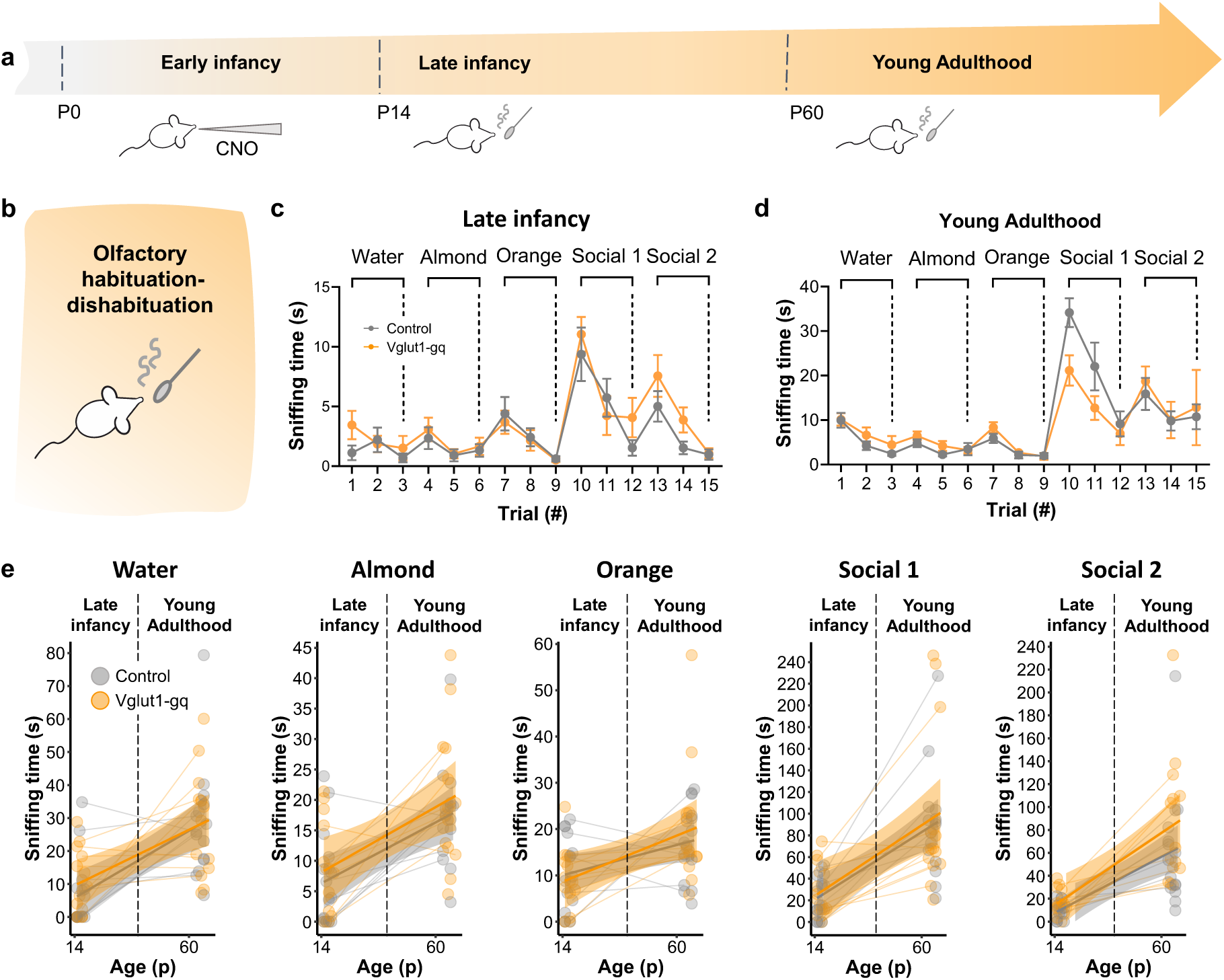
Developmental E:I imbalance does not alter olfactory discrimination. **(a)** Timeline of testing. **(b)** Schematic of the olfactory habituation-dishabituation task. **(c)** Sniffing time for water, almond, orange, and two unfamiliar social odors across trials in Vglut1-gq (n=14) and control mice (n=15) in late infancy and **(d)** young adulthood. **(e)** Quantification of sniffing time for each odor across ages. No significant group differences were observed for any odor (p>0.153, all odors).

**Supplementary Figure 4.**
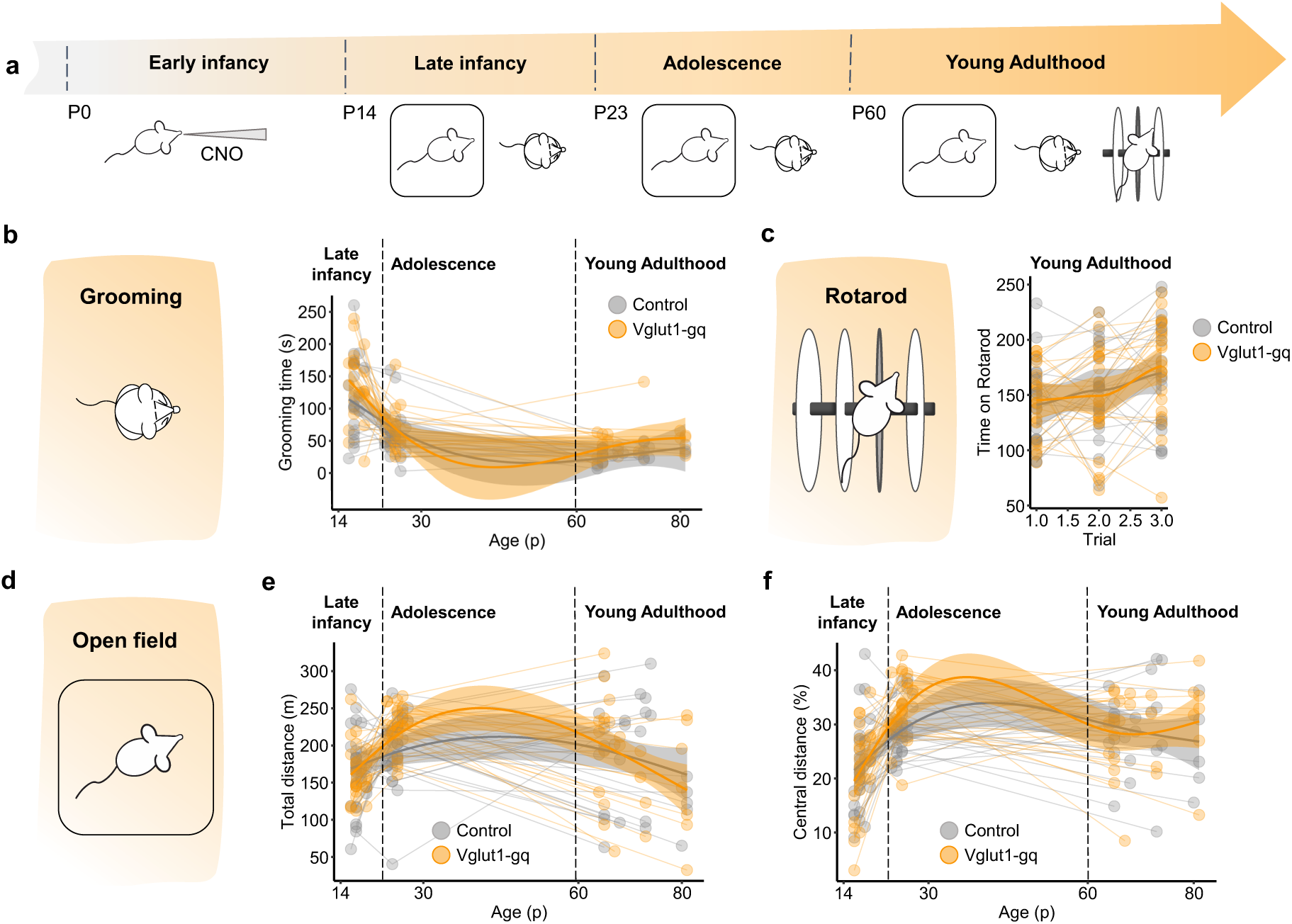
Grooming, rotarod and open field tests. **(a)** Timeline of testing. Open field measurements were carried out longitudinally in the same cohort of mice, which also underwent rotarod testing in adulthood. **(b)** Grooming time in Vglut1-gq (n=22) and control mice (n=20) across ages. **(c)** Time spent on the rotarod in adulthood (Vglut1-gq, n=23; control, n=20). **(d)** Schematic of the open field test. **(e)** Total distance travelled and **(f)** percentage of distance travelled in the center of the open field across ages (Vglut1-gq, n=26; control, n=27). Data are individual animals (dots). Shaded areas indicate mean ± 95% CI.

**Supplementary Figure 5.**
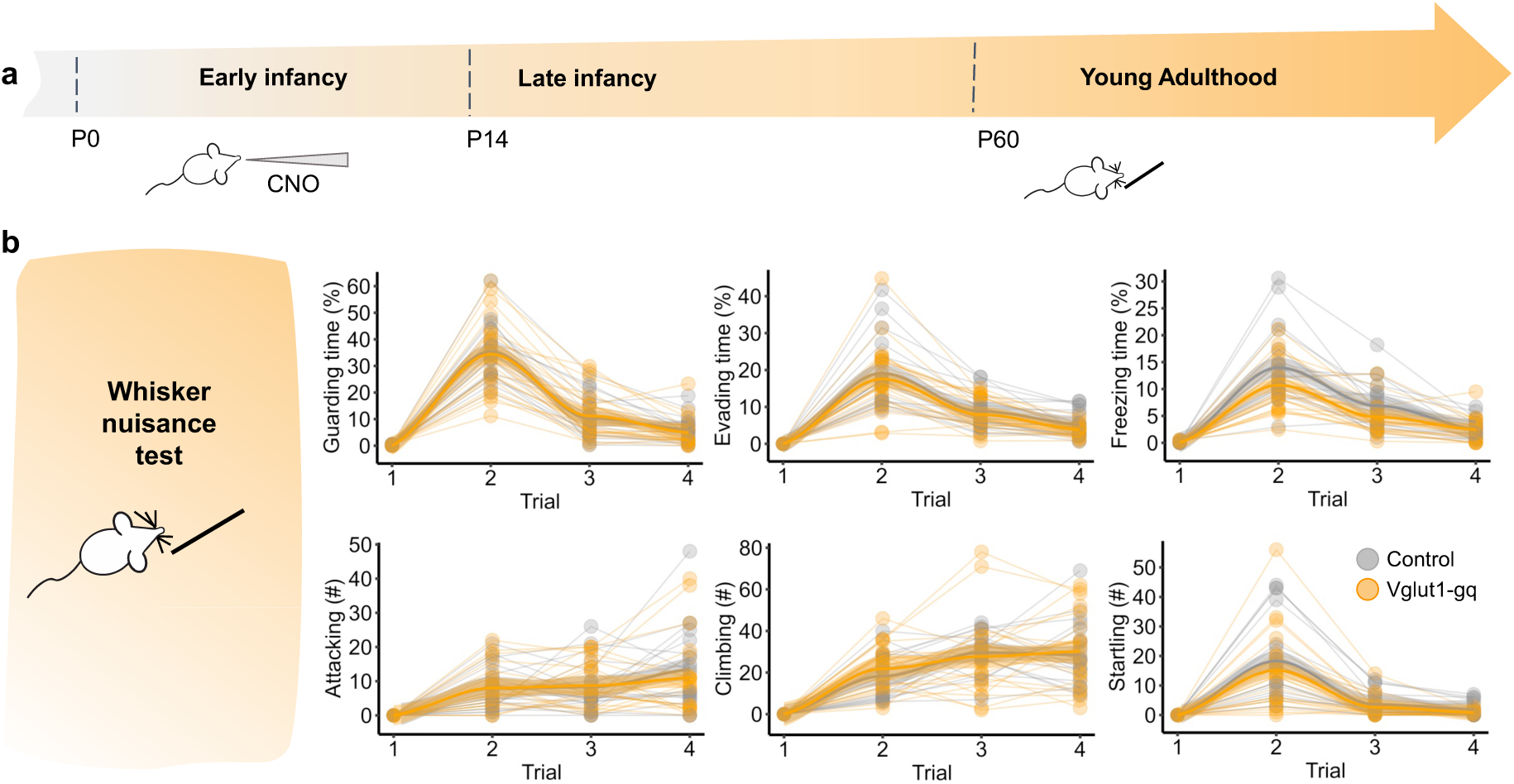
Vglut1-gq mice show preserved tactile responses. **(a)** Developmental timeline of testing. **(b)** Behavioral response to whisker stimulation across trials in adulthood. Vglut1-gq mice (n=23) and controls (n=20) showed comparable levels of evading, guarding, freezing, aiacking, climbing, and startling in response to mechanical whisker stimulation. Data are individual animals (dots); shaded areas show mean ± 95% CI.

**Supplementary Figure 6.**
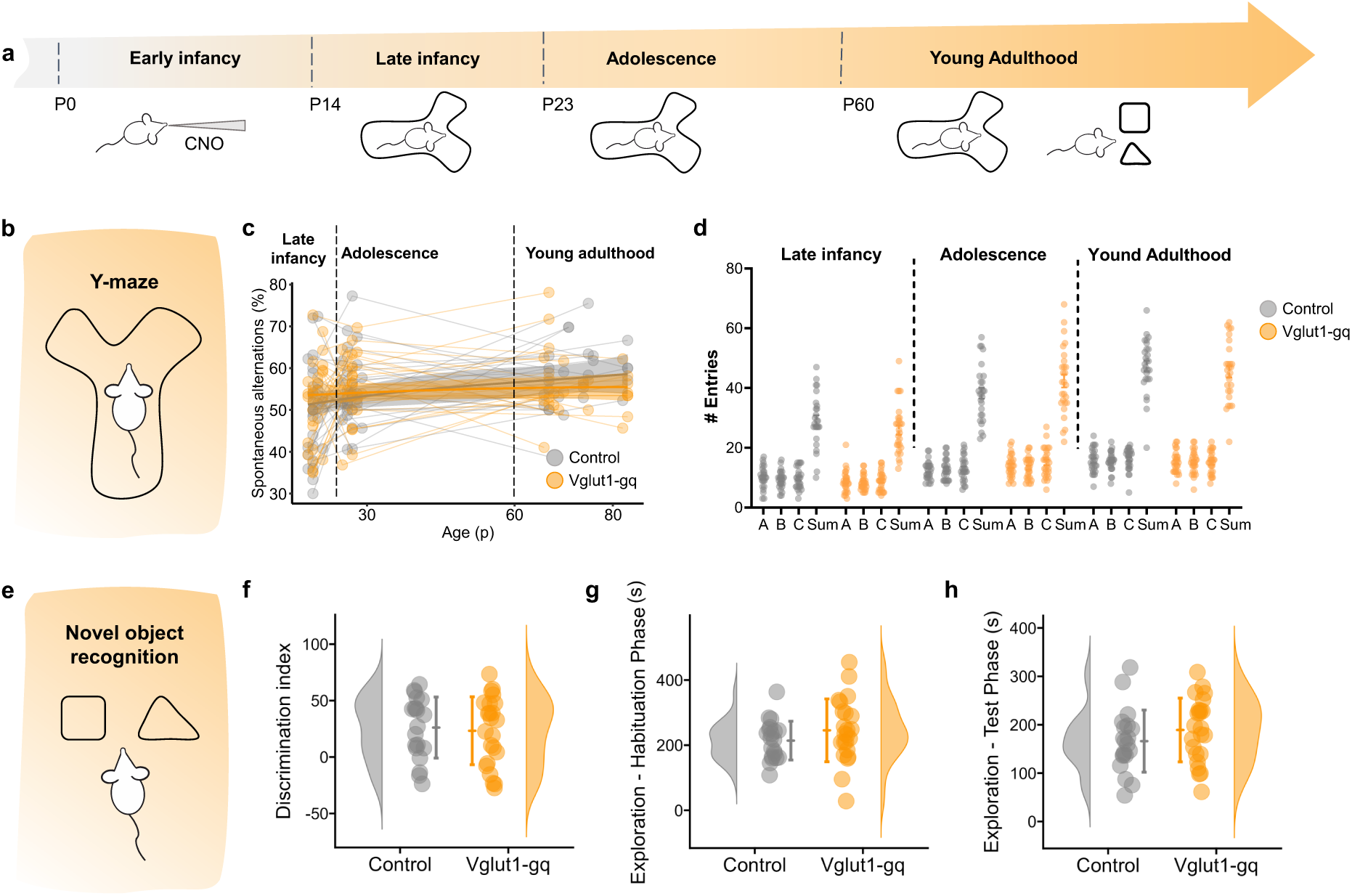
Y-maze and novel object recognition tests. **(a)** Timeline of behavioral tests. The Y-maze test was longitudinally carried out on the same cohort of mice, which also underwent novel object recognition testing in adulthood. **(b)** Schematic of the Y-maze test. **(c)** Percentage of spontaneous arm alternations across ages in Vglut1-gq (n=26) and control (n=27) mice. **(d)** Corresponding arm entries across ages (Sum=A + B +C). **(e)** Schematic of the novel object recognition test. **(f)** Discrimination index in adulthood (Vglut1-gq, n=23; control, n=20). **(g)** Total exploration time during the habituation phase. **(h)** Total Exploration time during the test phase. Data are individual animals (dots); shaded areas in (c) show mean ± 95% CI.

**Supplementary Figure 7.**
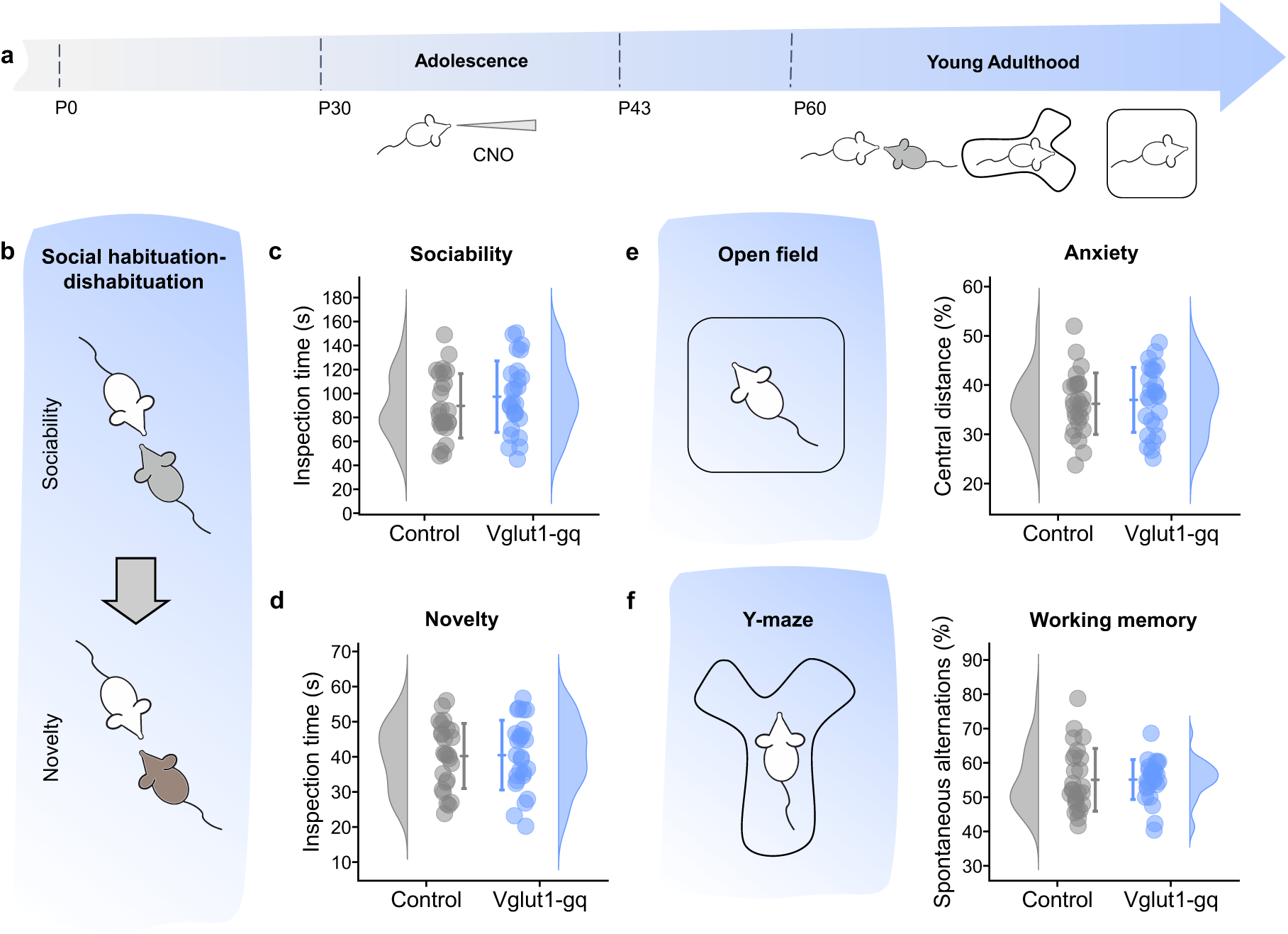
E:I imbalance during adolescence does not alter social behavior. **(a)** Experimental timeline. **(b)** Schematic of the social habituation/dishabituation test (Vglut1-gq n=26, control n=26). **(c)** Inspection time during habituation trials. **(d)** Inspection time during the novelty trial. **(e)** Percentage of distance travelled in the center of the open field arena (Vglut1-gq n=26, control n=26) mice. **(f)** Percentage of spontaneous arm alternations (Vglut1-gq n=26, control n=26). Data are individual animals (dots). Bars indicate mean ± SD.

**Supplementary Figure 8.**
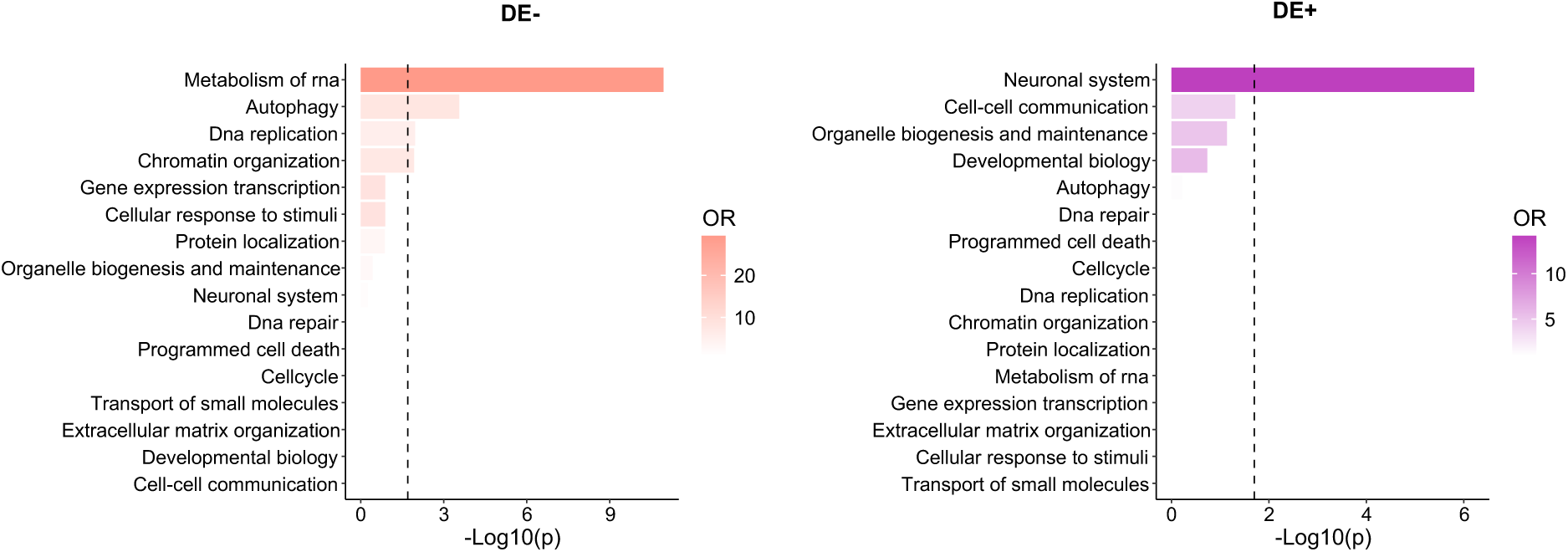
Pathway enrichment analysis for DE- and DE+ genes. Reactome pathway enrichment analysis for DE– (leT) and DE+ genes (right). The doied line marks significance at FDR q<0.05.

**Supplementary Figure 9.**
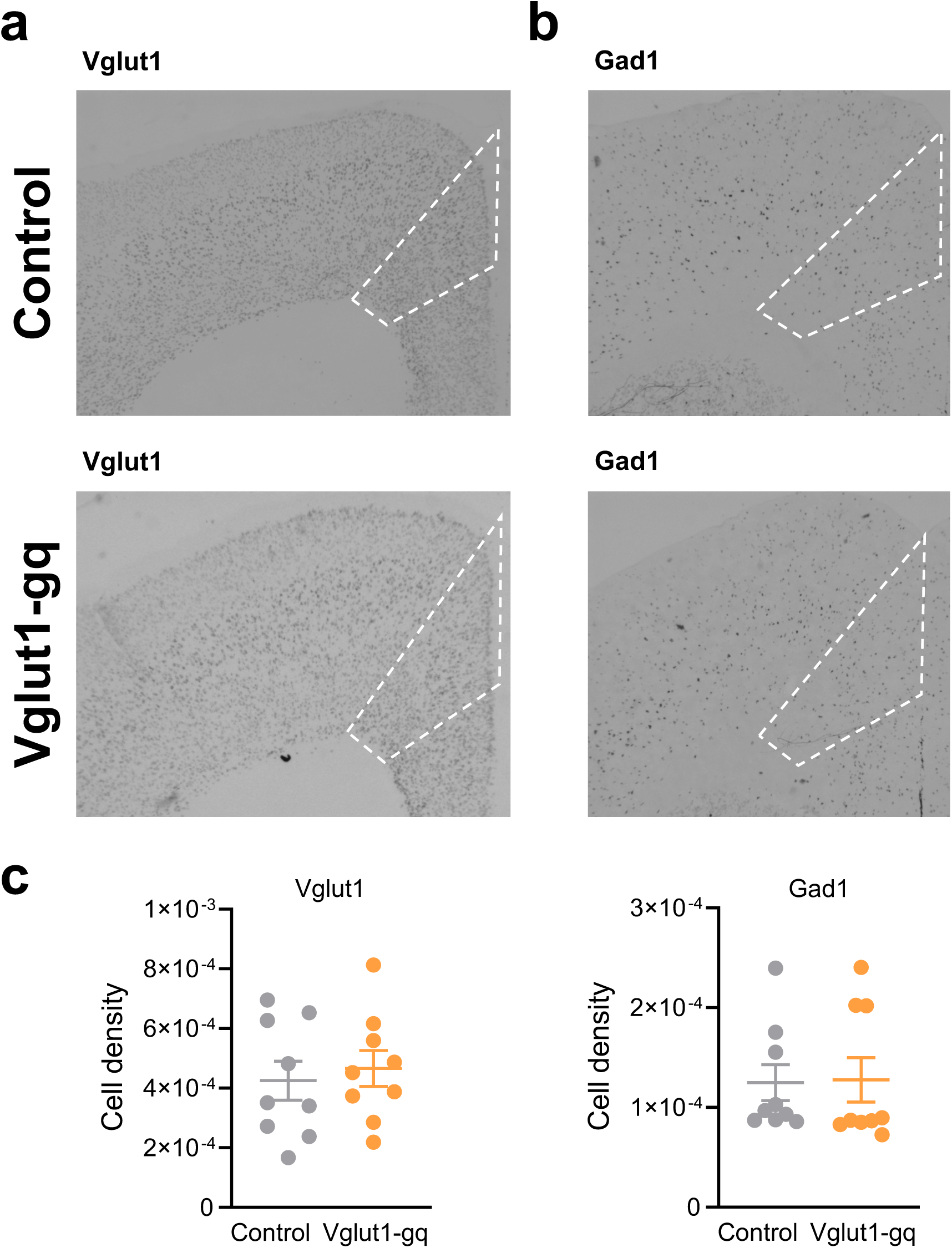
In situ hybridization of Vglut1 and Gad1 in adult mice. Representative ISH images for Vglut1 **(a)** and Gad1 **(b)** in the PFC of control (top) and Vglut1-gq (boiom) mice. **(c)** Quantification of cell density for Vglut1 (leT) and Gad1 (right) in control and Vglut1-gq mice. Data represents cell counts in three individual sections obtained in n=3 control and n=3 Vglut1-gq mice; bars show mean ± 95% CI.

## References

1. Shaw KA, et al. Prevalence and Early Identification of Autism Spectrum Disorder Among Children Aged 4 and 8 Years - Autism and Developmental Disabilities Monitoring Network, 16 Sites, United States, 2022. MMWR Surveill Summ 74, 1–22 (2025).

2. Lord C, et al. Autism spectrum disorder. Nature Reviews Disease Primers 6, 5 (2020).

3. Lombardo MV, Lai MC, Baron-Cohen S. Big data approaches to decomposing heterogeneity across the autism spectrum. Mol Psychiatry 24, 1435–1450 (2019).

4. de la Torre-Ubieta L, Won H, Stein JL, Geschwind DH. Advancing the understanding of autism disease mechanisms through genetics. Nat Med 22, 345–361 (2016).

5. Willsey HR, Willsey AJ, Wang B, State MW. Genomics, convergent neuroscience and progress in understanding autism spectrum disorder. Nat Rev Neurosci 23, 323–341 (2022).

6. Satterstrom FK, et al. Large-Scale Exome Sequencing Study Implicates Both Developmental and Functional Changes in the Neurobiology of Autism. Cell 180, 568–584 e523 (2020).

7. Trost B, et al. Genomic architecture of autism from comprehensive whole-genome sequence annotation. Cell 185, 4409–4427 e4418 (2022).

8. Meltzer A, Van de Water J. The Role of the Immune System in Autism Spectrum Disorder. Neuropsychopharmacology 42, 284–298 (2017).

9. Krumm N, O’Roak BJ, Shendure J, Eichler EE. A de novo convergence of autism genetics and molecular neuroscience. Trends Neurosci 37, 95–105 (2014).

10. Lombardo MV, Moon HM, Su J, Palmer TD, Courchesne E, Pramparo T. Maternal immune activation dysregulation of the fetal brain transcriptome and relevance to the pathophysiology of autism spectrum disorder. Mol Psychiatry 23, 1001–1013 (2018).

11. Rubenstein JLR, Merzenich MM. Model of autism: increased ratio of excitation/inhibition in key neural systems. *Genes*, Brain and Behavior 2, 255–267 (2003).

12. Nelson SB, Valakh V. Excitatory/Inhibitory Balance and Circuit Homeostasis in Autism Spectrum Disorders. Neuron 87, 684–698 (2015).

13. Bozzi Y, Provenzano G, Casarosa S. Neurobiological bases of autism-epilepsy comorbidity: a focus on excitation/inhibition imbalance. Eur J Neurosci 47, 534–548 (2018).

14. Bourgeron T. A synaptic trek to autism. Curr Opin Neurobiol 19, 231–234 (2009).

15. Penagarikano O, et al. Absence of CNTNAP2 leads to epilepsy, neuronal migration abnormalities, and core autism-related deficits. Cell 147, 235–246 (2011).

16. Peca J, et al. Shank3 mutant mice display autistic-like behaviours and striatal dysfunction. Nature 472, 437–442 (2011).

17. Canitano R. Epilepsy in autism spectrum disorders. Eur Child Adolesc Psychiatry 16, 61–66 (2007).

18. Lewine JD, et al. Magnetoencephalographic patterns of epileptiform activity in children with regressive autism spectrum disorders. Pediatrics 104, 405–418 (1999).

19. Tuchman R, Cuccaro M. Epilepsy and autism: neurodevelopmental perspective. Curr Neurol Neurosci Rep 11, 428–434 (2011).

20. Sohal VS, Rubenstein JLR. Excitation-inhibition balance as a framework for investigating mechanisms in neuropsychiatric disorders. Mol Psychiatry 24, 1248–1257 (2019).

21. Antoine MW, Langberg T, Schnepel P, Feldman DE. Increased Excitation-Inhibition Ratio Stabilizes Synapse and Circuit Excitability in Four Autism Mouse Models. Neuron 101, 648–661.e644 (2019).

22. Bertelsen N, et al. Electrophysiologically-defined excitation-inhibition autism neurosubtypes. medRxiv, 2023.2011.2022.23298729 (2024).

23. Boulting GL, et al. Activity-dependent regulome of human GABAergic neurons reveals new patterns of gene regulation and neurological disease heritability. Nature Neuroscience 24, 437–448 (2021).

24. Reh RK, et al. Critical period regulation across multiple timescales. Proceedings of the National Academy of Sciences 117, 23242–23251 (2020).

25. Alexander GM, et al. Remote control of neuronal activity in transgenic mice expressing evolved G protein-coupled receptors. Neuron 63, 27–39 (2009).

26. Minelli A, Edwards RH, Manzoni T, Conti F. Postnatal development of the glutamate vesicular transporter VGLUT1 in rat cerebral cortex. Developmental Brain Research 140, 309–314 (2003).

27. Harris JA, et al. Anatomical characterization of Cre driver mice for neural circuit mapping and manipulation. Front Neural Circuits 8, 76 (2014).

28. Liguz-Lecznar M, Skangiel-Kramska J. Vesicular glutamate transporters VGLUT1 and VGLUT2 in the developing mouse barrel cortex. International Journal of Developmental Neuroscience 25, 107–114 (2007).

29. Boulland J-L, et al. Expression of the vesicular glutamate transporters during development indicates the widespread corelease of multiple neurotransmitters. Journal of Comparative Neurology 480, 264–280 (2004).

30. Guma E, Plitman E, Chakravarty MM. The role of maternal immune activation in altering the neurodevelopmental trajectories of offspring: A translational review of neuroimaging studies with implications for autism spectrum disorder and schizophrenia. Neuroscience & Biobehavioral Reviews 104, 141–157 (2019).

31. Yang M, Crawley JN. Simple behavioral assessment of mouse olfaction. Curr Protoc Neurosci Chapter 8, Unit 8.24 (2009).

32. Silverman JL, Yang M, Lord C, Crawley JN. Behavioural phenotyping assays for mouse models of autism. Nat Rev Neurosci 11, 490–502 (2010).

33. Kraeuter AK, Guest PC, Sarnyai Z. The Open Field Test for Measuring Locomotor Activity and Anxiety-Like Behavior. Methods Mol Biol 1916, 99–103 (2019).

34. Deacon RM. Measuring motor coordination in mice. J Vis Exp, e2609 (2013).

35. Balasco L, Chelini G, Bozzi Y, Provenzano G. Whisker Nuisance Test: A Valuable Tool to Assess Tactile Hypersensitivity in Mice. Bio Protoc 9, e3331 (2019).

36. Nabel EM, et al. Adolescent frontal top-down neurons receive heightened local drive to establish adult attentional behavior in mice. Nature Communications 11, 3983 (2020).

37. Flavell SW, Greenberg ME. Signaling mechanisms linking neuronal activity to gene expression and plasticity of the nervous system. Annu Rev Neurosci 31, 563–590 (2008).

38. Ebert DH, Greenberg ME. Activity-dependent neuronal signalling and autism spectrum disorder. Nature 493, 327–337 (2013).

39. Gandal MJ, et al. Broad transcriptomic dysregulation occurs across the cerebral cortex in ASD. Nature 611, 532–539 (2022).

40. Yizhar O, et al. Neocortical excitation/inhibition balance in information processing and social dysfunction. Nature 477, 171–178 (2011).

41. Silverman JL, et al. GABAB Receptor Agonist R-Baclofen Reverses Social Deficits and Reduces Repetitive Behavior in Two Mouse Models of Autism. Neuropsychopharmacology 40, 2228–2239 (2015).

42. Savardi A, et al. Discovery of a Small Molecule Drug Candidate for Selective NKCC1 Inhibition in Brain Disorders. Chem 6, 2073–2096 (2020).

43. Alvino F, et al. Tracking the Developmental Trajectory of 22q11. 2 Deletion Syndrome in a Mouse Model. In: NEUROPSYCHOPHARMACOLOGY). SPRINGERNATURE CAMPUS, 4 CRINAN ST, LONDON, N1 9XW, ENGLAND (2021).

44. Pagani M, et al. Biological subtyping of autism via cross-species fMRI. bioRxiv, 2025.2003.2004.641400 (2025).

45. Pagani M, Gutierrez-Barragan D, de Guzman AE, Xu T, Gozzi A. Mapping and comparing fMRI connectivity networks across species. Commun Biol 6, 1238 (2023).

46. Whitesell JD, et al. Regional, Layer, and Cell-Type-Specific Connectivity of the Mouse Default Mode Network. Neuron 109, 545–559 e548 (2021).

47. Mars RB, Neubert FX, Noonan MP, Sallet J, Toni I, Rushworth MF. On the relationship between the “default mode network” and the “social brain“. Front Hum Neurosci 6, 189 (2012).

48. Ko J. Neuroanatomical Substrates of Rodent Social Behavior: The Medial Prefrontal Cortex and Its Projection Patterns. Frontiers in Neural Circuits 11, (2017).

49. McIntosh AR, Bookstein FL, Haxby JV, Grady CL. Spatial pattern analysis of functional brain images using partial least squares. Neuroimage 3, 143–157 (1996).

50. Vogel P, Hahn J, Duvarci S, Sigurdsson T. Prefrontal pyramidal neurons are critical for all phases of working memory. Cell Reports 39, 110659 (2022).

51. Adhikari A, et al. Basomedial amygdala mediates top-down control of anxiety and fear. Nature 527, 179–185 (2015).

52. Marek S, et al. Reproducible brain-wide association studies require thousands of individuals. Nature 603, 654–660 (2022).

53. O’Donnell C, Gonçalves JT, Portera-Cailliau C, Sejnowski TJ. Beyond excitation/inhibition imbalance in multidimensional models of neural circuit changes in brain disorders. eLife 6, e26724 (2017).

54. Pati S, et al. Chronic postnatal chemogenetic activation of forebrain excitatory neurons evokes persistent changes in mood behavior. eLife 9, e56171 (2020).

55. Bitzenhofer SH, Pöpplau JA, Chini M, Marquardt A, Hanganu-Opatz IL. A transient developmental increase in prefrontal activity alters network maturation and causes cognitive dysfunction in adult mice. Neuron 109, 1350–1364.e1356 (2021).

56. Zerbi V, et al. Brain mapping across 16 autism mouse models reveals a spectrum of functional connectivity subtypes. Mol Psychiatry 26, 7610–7620 (2021).

57. Hong SJ, et al. Toward Neurosubtypes in Autism. Biol Psychiatry 88, 111–128 (2020).

58. Buch AM, Vértes PE, Seidlitz J, Kim SH, Grosenick L, Liston C. Molecular and network-level mechanisms explaining individual differences in autism spectrum disorder. Nature Neuroscience 26, 650–663 (2023).

59. Lombardo MV, et al. Large-scale associations between the leukocyte transcriptome and BOLD responses to speech differ in autism early language outcome subtypes. Nature Neuroscience 21, 1680–1688 (2018).

60. Vicari S, et al. Copy number variants in autism spectrum disorders. Progress in Neuro-Psychopharmacology and Biological Psychiatry 92, 421–427 (2019).

61. Auerbach BD, Osterweil EK, Bear MF. Mutations causing syndromic autism define an axis of synaptic pathophysiology. Nature 480, 63–68 (2011).

62. Oostra BA, Willemsen R. A fragile balance: FMR1 expression levels. Human Molecular Genetics 12, R249–R257 (2003).

63. Xu M, et al. Aberrant brain functional and structural developments in MECP2 duplication rats. Neurobiol Dis 173, 105838 (2022).

64. Pinto D, et al. Functional impact of global rare copy number variation in autism spectrum disorders. Nature 466, 368–372 (2010).

65. Sanders SJ, et al. De novo mutations revealed by whole-exome sequencing are strongly associated with autism. Nature 485, 237–241 (2012).

66. Han K, et al. SHANK3 overexpression causes manic-like behaviour with unique pharmacogenetic properties. Nature 503, 72–77 (2013).

67. Qin L, Ma K, Yan Z. Chemogenetic Activation of Prefrontal Cortex in Shank3-Deficient Mice Ameliorates Social Deficits, NMDAR Hypofunction, and Sgk2 Downregulation. iScience 17, 24–35 (2019).

68. Wang W, et al. Chemogenetic Activation of Prefrontal Cortex Rescues Synaptic and Behavioral Deficits in a Mouse Model of 16p11.2 Deletion Syndrome. J Neurosci 38, 5939–5948 (2018).

69. Sastre-Yague D, et al. Cortical excitatory-inhibitory balance critically biases brain-wide fMRI connectivity in preparation, (2025).

70. Rocchi F, et al. Increased fMRI connectivity upon chemogenetic inhibition of the mouse prefrontal cortex. Nature Communications 13, 1056 (2022).

71. Dautan D, et al. Cortico-cortical transfer of socially derived information gates emotion recognition. Nature Neuroscience 27, 1318–1332 (2024).

72. Chen Z, et al. A prefrontal-thalamic circuit encodes social information for social recognition. Nat Commun 15, 1036 (2024).

73. Park G, et al. Social isolation impairs the prefrontal-nucleus accumbens circuit subserving social recognition in mice. Cell Rep 35, 109104 (2021).

74. Giorgi A, et al. Brain-wide Mapping of Endogenous Serotonergic Transmission via Chemogenetic fMRI. Cell Reports 21, 910–918 (2017).

75. Migliarini S, Pacini G, Pelosi B, Lunardi G, Pasqualetti M. Lack of brain serotonin affects postnatal development and serotonergic neuronal circuitry formation. Mol Psychiatry 18, 1106–1118 (2013).

76. Gomez JL, et al. Chemogenetics revealed: DREADD occupancy and activation via converted clozapine. Science 357, 503–507 (2017).

77. Jendryka M, et al. Pharmacokinetic and pharmacodynamic actions of clozapine-N-oxide, clozapine, and compound 21 in DREADD-based chemogenetics in mice. Scientific Reports 9, 4522 (2019).

78. Teissier A, et al. Activity of Raphé Serotonergic Neurons Controls Emotional Behaviors. Cell Rep 13, 1965–1976 (2015).

79. Bicks LK, et al. Prefrontal parvalbumin interneurons require juvenile social experience to establish adult social behavior. Nat Commun 11, 1003 (2020).

80. Chao OY, et al. Targeting inhibitory cerebellar circuitry to alleviate behavioral deficits in a mouse model for studying idiopathic autism. Neuropsychopharmacology 45, 1159–1170 (2020).

81. Kayyal H, Yiannakas A, Kolatt Chandran S, Khamaisy M, Sharma V, Rosenblum K. Activity of Insula to Basolateral Amygdala Projecting Neurons is Necessary and Sufficient for Taste Valence Representation. J Neurosci 39, 9369–9382 (2019).

82. Quiroga RQ, Nadasdy Z, Ben-Shaul Y. Unsupervised spike detection and sorting with wavelets and superparamagnetic clustering. Neural Comput 16, 1661–1687 (2004).

83. Welch P. The use of fast Fourier transform for the estimation of power spectra: A method based on time averaging over short, modified periodograms. IEEE Transactions on Audio and Electroacoustics 15, 70–73 (1967).

84. Belitski A, et al. Low-Frequency Local Field Potentials and Spikes in Primary Visual Cortex Convey Independent Visual Information. The Journal of Neuroscience 28, 5696–5709 (2008).

85. Maris E, Oostenveld R. Nonparametric statistical testing of EEG- and MEG-data. J Neurosci Methods 164, 177–190 (2007).

86. Ferguson JN, Young LJ, Hearn EF, Matzuk MM, Insel TR, Winslow JT. Social amnesia in mice lacking the oxytocin gene. Nature Genetics 25, 284–288 (2000).

87. Ferretti V, et al. Oxytocin Signaling in the Central Amygdala Modulates Emotion Discrimination in Mice. Curr Biol 29, 1938–1953.e1936 (2019).

88. Rein B, Ma K, Yan Z. A standardized social preference protocol for measuring social deficits in mouse models of autism. Nature Protocols 15, 3464–3477 (2020).

89. McFarlane HG, Kusek GK, Yang M, Phoenix JL, Bolivar VJ, Crawley JN. Autism-like behavioral phenotypes in BTBR T+tf/J mice. *Genes*, Brain and Behavior 7, 152–163 (2008).

90. Faizi M, et al. Comprehensive behavioral phenotyping of Ts65Dn mouse model of Down syndrome: activation of β1-adrenergic receptor by xamoterol as a potential cognitive enhancer. Neurobiol Dis 43, 397–413 (2011).

91. Lueptow LM. Novel Object Recognition Test for the Investigation of Learning and Memory in Mice. J Vis Exp, (2017).

92. Montani C, et al. Sex-biasing influence of autism-associated Ube3a gene overdosage at connectomic, behavioral and transcriptomic levels. bioRxiv, 2022.2010.2025.513747 (2022).

93. Robinson MD, Oshlack A. A scaling normalization method for differential expression analysis of RNA-seq data. Genome Biology 11, R25 (2010).

94. Law CW, Chen Y, Shi W, Smyth GK. voom: precision weights unlock linear model analysis tools for RNA-seq read counts. Genome Biology 15, R29 (2014).

95. Chen EY, et al. Enrichr: interactive and collaborative HTML5 gene list enrichment analysis tool. BMC Bioinformatics 14, 128 (2013).

96. Kuleshov MV, et al. Enrichr: a comprehensive gene set enrichment analysis web server 2016 update. Nucleic Acids Res 44, W90–97 (2016).

97. Gandal MJ, et al. Transcriptome-wide isoform-level dysregulation in ASD, schizophrenia, and bipolar disorder. Science 362, eaat8127 (2018).

98. Wilfert AB, et al. Recent ultra-rare inherited variants implicate new autism candidate risk genes. Nat Genet 53, 1125–1134 (2021).

99. Grove J, et al. Identification of common genetic risk variants for autism spectrum disorder. Nat Genet 51, 431–444 (2019).

100. Koopmans F, et al. SynGO: An Evidence-Based, Expert-Curated Knowledge Base for the Synapse. Neuron 103, 217–234.e214 (2019).

101. Niell CM, Stryker MP. Highly Selective Receptive Fields in Mouse Visual Cortex. The Journal of Neuroscience 28, 7520–7536 (2008).

102. Dana H, et al. Thy1-GCaMP6 transgenic mice for neuronal population imaging in vivo. PLoS One 9, e108697 (2014).

103. Chornyy S, et al. Cellular-resolution monitoring of ischemic stroke pathologies in the rat cortex. Biomed Opt Express 12, 4901–4919 (2021).

104. Zhuang J, et al. An extended retinotopic map of mouse cortex. eLife 6, e18372 (2017).

105. Gutierrez-Barragan D, Basson MA, Panzeri S, Gozzi A. Infraslow State Fluctuations Govern Spontaneous fMRI Network Dynamics. Curr Biol 29, 2295–2306.e2295 (2019).

106. Sforazzini F, Schwarz AJ, Galbusera A, Bifone A, Gozzi A. Distributed BOLD and CBV-weighted resting-state networks in the mouse brain. Neuroimage 87, 403–415 (2014).

107. Gutierrez-Barragan D, et al. Unique spatiotemporal fMRI dynamics in the awake mouse brain. Curr Biol 32, 631–644 e636 (2022).

108. Liska A, Galbusera A, Schwarz AJ, Gozzi A. Functional connectivity hubs of the mouse brain. Neuroimage 115, 281–291 (2015).

109. Gorgolewski KJ, et al. Tight fitting genes: finding relations between statistical maps and gene expression patterns. F1000Posters 5, 10.7490 (2014).

110. Rubinov M, Ypma RJ, Watson C, Bullmore ET. Wiring cost and topological participation of the mouse brain connectome. Proceedings of the National Academy of Sciences 112, 10032–10037 (2015).

111. Markello RD, Arnatkeviciute A, Poline J-B, Fulcher BD, Fornito A, Misic B. Standardizing workflows in imaging transcriptomics with the abagen toolbox. elife 10, e72129 (2021).

112. Kebets V, et al. Somatosensory-Motor Dysconnectivity Spans Multiple Transdiagnostic Dimensions of Psychopathology. Biological Psychiatry 86, 779–791 (2019).

113. Krishnan A, Williams LJ, McIntosh AR, Abdi H. Partial Least Squares (PLS) methods for neuroimaging: a tutorial and review. Neuroimage 56, 455–475 (2011).

